# Sensor placement causes outcome-dependent bias in ambulatory light-exposure estimates

**DOI:** 10.64898/2026.07.28.741277

**Authors:** Johannes Zauner, Sietse W. de Vries, Altug Didikoglu, Juliëtte van Duijnhoven, Manuel Spitschan

**Author notes:** These authors contributed equally to this work (Johannes Zauner, Sietse W. de Vries). Correspondence: Johannes Zauner* < >, Sietse W. de Vries* < >.

## Abstract

Population studies use wearable sensors to investigate light as an environmental determinant of health, yet anatomical placement may misclassify exposure across outcomes. We analysed concurrent free-living measurements from one dosimeter model worn near the eyes, chest, and wrist across seven countries. Generalised additive mixed models characterised momentary errors; mixed models compared 54 daily metrics; hierarchical bootstrap estimated validation precision. Context-specific eye-level underestimation ranged from −23.1% to −4.2% at the chest and −54.7% to −30.3% at the wrist. Across 54 metrics, median absolute bias was 1% at the chest and 2% at the wrist; timing outcomes were robust, whereas level and temporal-dynamics outcomes were more placement-sensitive, with maximum absolute biases of 50% and 84%, respectively. With seven days, most metric classes reached 5% precision with 2–16 participants; temporal dynamics required 33 at the chest and 59 at the wrist. Sensor placement should match the intended outcome, with validation subsamples supporting scalable exposure epidemiology.

## Introduction

Ocular light is a physical environmental exposure with effects extending beyond vision. Light reaching the eye regulates circadian entrainment, sleep–wake timing, acute alertness and other physiological and behavioural responses^1–4^. Emerging epidemiological evidence further associates patterns of daytime and nighttime light exposure with sleep, mental health, metabolic outcomes, and mortality^5–9^. These findings have increased interest in personal light exposure as a potentially modifiable component of the human exposome^4^. However, evaluating its health relevance depends on accurately characterising the light that individuals receive as they move through natural and built environments.

Wearable dosimeters (also called light loggers) make it possible to capture this temporally dynamic exposure under free-living conditions^10^. Unlike stationary measurements or estimates based on outdoor conditions, they accompany participants across locations and activities and can record exposure patterns arising from the interaction of environmental light, architecture and individual behaviour^11–14^. This has supported increasingly large observational studies, such as the UK Biobank^6–9^, and creates opportunities to integrate personal light exposure into environmental epidemiology. At the same time, wearable assessment introduces a fundamental problem: the dosimeter is rarely positioned at the anatomical site corresponding to the exposure of interest, the eye.

For research on non-visual responses to light, the target quantity is the spectrally resolved optical radiation incident at the retina, which cannot be measured directly^15^. Exposure is therefore ideally measured close to the corneal plane, with a sensor orientation approximating the participant’s direction of view^14–17^. Near-eye devices can, however, be conspicuous or uncomfortable and may limit adherence and study duration^18^. Dosimeters placed on the chest or wrist impose less participant burden^18^ and are consequently attractive for larger and longer studies^16^. The choice of placement (sometimes termed wearing position) thus represents a trade-off between measurement fidelity and operational scalability. Whether it is acceptable depends not only on the average agreement between placements, but on how placement-related differences vary across participants, environments and activities and how they affect the summary metrics used in subsequent analyses. Throughout this manuscript, *eye-level*, *near-corneal*, and *glasses-level* refer to the same spectacle-mounted reference position, whereas *body-worn* refers collectively to the chest and wrist placements. We use *glasses*, *chest*, and *wrist* when referring to the three specific sensor placements.

The influence of anatomical placement has long been recognised in optical-radiation dosimetry, first in research on ultraviolet radiation (UVR), where studies quantified relative UVR doses across body surfaces using a mannequin^19^, characterised surface-specific doses across occupations and recreational activities^20^, and assessed whether wrist-worn dosimeters correlate with exposure on the top of the head^21^. Comparable concerns later emerged in personal visible-light dosimetry, where field studies compared measurements at the eye, chest and wrist^22–28^ (Supplementary Table S1). These protocols are broadly similar, differing mainly in participant number (one to twenty-nine), duration (hours to days), dosimeter type, and participant occupation and location. Beyond raw values (e.g., photopic illuminances), some examined errors in derived metrics such as circadian stimulus^24^, luminous exposure^26^, and time above threshold^25,27^. Collectively, they suggest that chest measurements approximate eye-level exposure more closely than wrist measurements, but report substantial variability and inconsistent directions of bias: depending on protocol and context, body-worn dosimeters have been found to either under- or overestimate near-eye measurements (see^29^).

Such inconsistencies are plausible because placement bias arises from the spatial structure of the surrounding illumination and from the participant’s posture and orientation. A chest-mounted dosimeter may be shaded by clothing, directed away from a light source, or exposed to light that does not reach the eyes. Wrist measurements are additionally affected by frequent changes in arm position and occlusion, e.g., from sleeves. Laboratory and simulation studies confirm that illuminance ratios between anatomical sites depend on source direction, light distribution, head and body orientation, and body morphology^29–33^. These establish the mechanisms through which placement affects validity, but controlled scenes cannot reproduce the diversity and temporal sequencing of everyday environments.

The practical consequences of placement bias also depend on the analytical scale and intended use of the data. For time-resolved analyses, differences between simultaneous measurements indicate how closely a proxy reproduces momentary eye-level exposure, and positive and negative deviations remain distinct. Epidemiological and intervention studies, however, commonly reduce light time series to daily or phase-specific metrics such as mean exposure, cumulative exposure, time above threshold, or the timing of exposure^34–36^. Under aggregation, overestimation during some periods can offset underestimation during others: a placement may poorly reproduce individual measurements yet yield a similar daily summary, or small systematic differences may accumulate into substantial metric differences. Correlation between placements alone therefore cannot establish fitness for a particular purpose. This is especially relevant to environmental epidemiology, where placement-dependent error may alter exposure ranking, attenuate associations, bias threshold-based classifications, or affect estimates of compliance with exposure recommendations.

Here, we evaluate placement bias using data from the MeLiDos project, a harmonised field study of wearable light exposure conducted across sites in Ghana, Costa Rica, Türkiye, Spain, Germany, the Netherlands and Sweden^11,13,37^. Participants wore calibrated dosimeters at the glasses, chest, and wrist placements during normal daily activities, with the glasses-mounted dosimeter positioned near the corneal plane and with repeated contextual reports. Using glasses-level measurements as the reference estimate of ocular exposure, we assess chest- and wrist-based proxies at two complementary levels. At the measurement level, we quantify time-resolved differences and how they vary with activity, primary light source, photoperiod, and indoor–outdoor context. At the outcome (metric) level, we determine how placement affects 54 commonly used daily and phase-specific exposure summaries. By evaluating both instantaneous fidelity and outcome-level agreement under diverse free-living conditions, we aim to establish when less burdensome placements^18^ provide fit-for-purpose estimates and when they may introduce consequential exposure misclassification.

## Methods

### Dataset

Data were obtained from the field-study component of the MeLiDos project (Metrology for wearable light loggers and optical radiation dosimeters)^13^. The study used a harmonised protocol across nine study sites in seven countries: Ghana, Costa Rica, Türkiye, Spain, Germany, the Netherlands, and Sweden^37^. Eight of these sites contain data for more than one simultaneous sensor placement. Tübingen (Germany), as the pilot site for the protocol, only used the glasses placement, and a different device type on the wrist. Thus, this ninth site will be disregarded in this study. The complete MeLiDos field-study dataset comprises 191 participants and includes temporally resolved personal light-exposure measurements, participant wear and sleep logs, repeated contextual assessments, and questionnaire data^38^. The data are permissively licensed (CC-BY4.0). See^11^ for details on device availability and measurement completeness between sites and participants.

We used recordings from participants with concurrent measurements at the glasses, chest, and wrist placements, together with the corresponding wear logs and hourly contextual reports. The glasses placement, mounted on spectacle frames near the corneal plane, served as the eye-level reference; chest and wrist were evaluated as the alternative body-worn placements. Contextual data comprised the participant’s reported activity (i.e., being awake at home, sleeping in bed, on the road with public transport/car, on the road with bike/on foot, working in the office/from home, working outdoors - including lunch break outdoors, free time outdoors, other) and the primary light source in the environment (electric light source indoors, electric light source outdoors, daylight indoors, daylight outdoors - including shade, emissive display light, darkness during sleep, light entering from outside during sleep). Inclusion criteria and processing steps are described below.

Light exposure was recorded with spectrally sensitive ActLumus devices (Condor Instruments, São Paulo, Brazil)^10,39^, sampling at 10-s intervals and using eight visible-range spectral channels to derive photopic and α-opic quantities, including melanopic equivalent daylight illuminance (melEDI)^40^. Devices were factory calibrated. Participants wore separate devices mounted centrally on non-prescription spectacle frames near the corneal plane, as a pendant at chest level, and with a wristband on the wrist. ActLumus devices have been independently evaluated against a criterion spectrometer under indoor electric LED and outdoor daylight conditions, showing a broad detection range (100% of the tested 2-100k lx), low inter-device variability, and comparatively high photopic accuracy^39^. Further details of the devices, calibration, wearing protocol, and field procedures are reported in the companion study^11^ and in Burcu et al.^41^.

### Software Resources

All data processing, statistical analyses, and visualisations were performed in R version 4.5.0^42^. Harmonised data from the individual MeLiDos study sites^38^ were accessed using *melidosData* version 1.0.6^43^. Personal light-exposure time series were processed, temporally aligned, annotated, and summarised using *LightLogR* version 0.10.0^36^. General data manipulation and visualisation were performed with packages from the *tidyverse* version 2.0.0^44^.

Generalized additive mixed models were fitted using *mgcv* version 1.9-4^45^, and model-derived smooths, differences, and predictions were processed using *gratia* version 0.11.2^46^. Linear and generalized linear mixed-effects models were fitted using *lme4* version 2.0-1^47^ and *glmmTMB* version 1.1.14^48^. Estimated marginal means and contrasts were calculated using *emmeans* version 2.0.3^49^, and model diagnostics and variance-explanation statistics were obtained using *performance* version 0.16.0^50^.

All analyses used scripted workflows for reproducibility. The analysis code, computational environment, and package-version information are available in a version-controlled repository https://github.com/tscnlab/ZaunerDeVriesEtAl_bioRxiv_2026 and an archived release (https://doi.org/10.5281/zenodo.21533915).

### Preprocessing and cleaning of data

The site-specific datasets provided by melidosData^43^ had already undergone several harmonisation steps. Raw dosimeter files were imported and each participant’s time series trimmed to the official trial dates in the study metadata. Implausible values attributable to device malfunction were removed per participant (e.g. constant values or strong, intermittent values around 10^4 lx throughout the day and night). Irregular sampling intervals and implicit gaps were resolved by regularising every series to a uniform 10-s epoch: values falling within a 10-s window were averaged, and a single measurement was assigned to the nearest boundary. The result was a continuous time series for each participant and placement, with explicit gaps at missing time-series instances^43^.

The target metric, and basis for the summary metrics, is melEDI as defined by CIE S026^40^, derived from the devices’ spectral sensors by device-internal computation.

Because the two analyses place different demands on the data, two preprocessing pipelines were applied downstream (Figure 1). The measurement-level analysis requires strict temporal alignment of concurrent observations but tolerates incomplete days, whereas the outcome-level analysis computes daily and multi-day summaries and therefore requires sufficiently complete days. Detailed filtering steps for each pipeline are described below.

**Figure 1:**
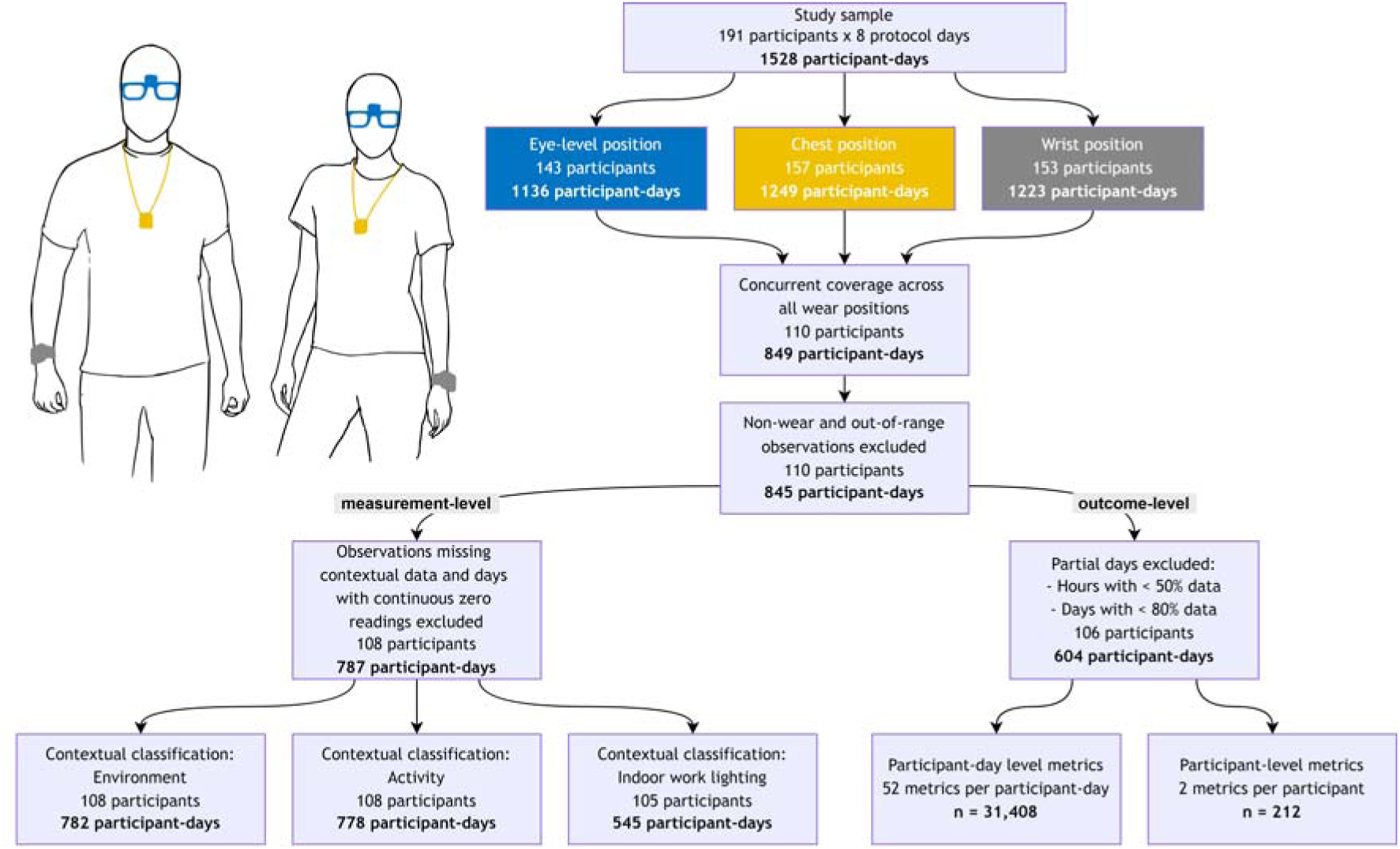
Data processing pipelines for measurement-level and outcome-level analyses.

Both analyses required log-transformation. Because light-exposure data frequently contain zeros - e.g., during complete darkness - for which the logarithm is undefined, a zero-inflated log_10_ transformation^51^ was applied using the *log_zero_inflated* function of *LightLogR*:

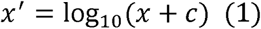

Where *c* = 0.1 lx melEDI is a small offset added before transformation. This maps 0 lx to *x′* = − 1 while preserving interpretability of the log_10_ scale for positive values. The offset lies below the dosimeters’ operating range and has no meaningful impact on results. The transformation is invertible via *x* = 10*^x^*^′^ − *c*. The metrics and responses to which it was applied are listed in Supplementary Table S2.

### Measurement-level analysis

melEDI recordings from the three dosimeters were temporally aligned and filtered as follows. Only time points with concurrent data from all three dosimeters were retained; observations during sleep and other documented non-wear periods (from the wear logs) were removed; only observations below the maximum operating range (<100,000 lx melEDI, per the manufacturer) were kept; and days on which all three dosimeters recorded continuous zeros were removed. Each remaining observation was paired with hourly contextual data - the activity performed and the primary light source, categorised per the study protocol^37^ - and observations lacking these entries were removed. The photoperiod state (day or night, from civil dawn and dusk) was matched to each observation. Retained participants and participant-days after each step are shown in Figure 1.

Measurements were log-transformed (Equation 1), and for each observation the placement error was calculated as the difference between a body-worn dosimeter’s measurement and the glasses reference. These momentary errors were aggregated to hourly resolution to match the contextual data. The hourly aggregated placement error *D_x_* is defined in Equation 2:

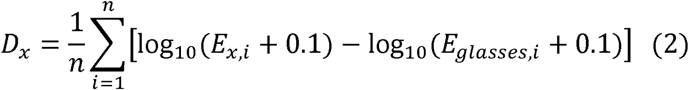

where *E_glasses,i_* is the glasses measurement and *E_x,i_* the measurement at placement *x*, both at timestamp *i*, and *n* is the number of measurements within the hour. A positive placement error indicates that the body-worn dosimeter overestimates glasses-level exposure.

### Outcome-level analysis

melEDI and photopic illuminance recordings from the glasses, chest, and wrist were prepared as above, yielding a single time series with concurrent measurements from all three placements. Two deviations applied: no filtering by missing activity or light-source entries, and sleep-time measurements were retained, as some daily metrics require sufficiently complete daily coverage. Two completeness filters were then applied: hours with more than 50% of expected 10-s observations missing were discarded, and days retaining less than 80% of their expected hourly bins were removed.

A comprehensive set of daily light-exposure metrics was computed per participant, day, and placement using *LightLogR* functions: duration above threshold (10, 250, and 1,000 lx melEDI), mean period above threshold, pulse analysis, brightest and darkest 10-hour period characteristics, timing of first and last threshold crossing, frequency of threshold crossings, the Barroso lighting metrics, centroid of light exposure, disparity index, midpoint of cumulative exposure, mean log-transformed melEDI, luminous dose, the melanopic daylight equivalent ratio (MDER), and the non-visual relative dose (nvRD). Interdaily stability and intradaily variability, which require multiple days, were computed per participant and placement across all available days.

Finally, two nighttime timing metrics (darkest 10-hour midpoint and onset) were rescaled to avoid circularity around midnight: values after 12:00 were shifted by −24 hours, representing late-evening times as negative values relative to midnight. This allows modelling with standard linear methods without midnight-boundary artefacts.

For metric definitions, see Hartmeyer and Andersen (2023)^35^, which also define metric classes, or categories, that will be used throughout this study: metrics based on duration, timing, spectrum, exposure history and temporal dynamics.

### Statistical analysis

No a priori power analysis was performed, as the sample size was fixed by the available dataset^38^. The analyses were primarily estimation-focused: effect estimates and confidence intervals characterised the magnitude, direction, and consistency of placement-dependent differences across contexts and metrics. Inferential tests were treated as aids rather than confirmatory hypothesis tests, and no adjustment for multiple comparisons was applied. Individual p-values should therefore not be interpreted in isolation; conclusions rest principally on effect sizes, uncertainty intervals, and patterns across related outcomes.

### Measurement-level analysis

Placement errors are influenced by both the surrounding illumination environment and the participant’s posture^29–31,33^. As illumination fields and postures were not directly quantified, four sets of contextual classifications (Supplementary Table S3) were defined as proxies: 1) environment (indoors or outdoors) × photoperiod (day or night); 2) environment × local time of day; 3) activity × photoperiod; and 4) the primary indoor-work light source. Activity categories were being awake at home, working (at home or in the office), travelling in a vehicle, and time outdoors; the last grouped walking or cycling, outdoor work, and outdoor leisure, each too sparsely reported to examine separately. Classifications 1) and 2) use data from the environment context (108 participants, 782 participant-days), 3) from the activity context (108 participant, 778 participant-days), and 4) from the indoor-work lighting context (105 participants, 545 participant-days). Number or participants and participant-days per set are shown in Figure 1.

Generalised additive mixed models (GAMMs)^52–54^ were fitted to model placement error as a function of these classifications. To avoid multicollinearity and aid interpretation, separate models were fitted for each set and each placement (chest and wrist), giving eight models. Classifications were fixed effects, except local time of day, which was modelled as a cyclic smooth. Because placement errors were heavy-tailed, a scaled t-distribution was specified for residuals. Random intercepts were specified for site, participant (nested within site), and day (nested within participant), reflecting the study protocol^37^. Although some explained little variance, all were retained as theoretically justified. A first-order autoregressive error structure (*AR* (1)) per participant accounted for temporal autocorrelation; its correlation parameter (*rho*) was set to the lag-1 residual autocorrelation from an initial model fitted without *AR*(1). Model formulas are given in Supplementary Table S4.

Models were fitted with the *bam* function (*mgcv* package) using fast REML (fREML) and discretisation, which provides benefits for computational speed for large datasets and an integrated autocorrelation correction for lag 1. Fixed-effect estimates agreed to three decimal places (maximum absolute difference < 0.001 log_10_ units) with the *gam* function using REML without discretisation (both without *AR*(1), indicating no meaningful impact.

All models converged. Because placement errors are differences between paired dosimeter measurements and the classifications are coarse descriptions of context, the deviance explained was modest (11.2%–32.8%). As the models were intended to identify systematic differences across contexts rather than to predict individual errors, this was considered acceptable. The degrees-of-freedom parameter (ν) of the scaled-t residuals was estimated at its lower bound (3) in all models, suggesting the true residuals may be heavier-tailed than the models can represent; consistent with this, residual distributions agreed fairly in the centre but deviated slightly in the tails (diagnostics are available in the repository). As no *mgcv* residual family accommodated heavier tails and further data transformation was inappropriate given the existing log transform, no further optimisation was feasible. Interpretation therefore rests primarily on estimated mean effects across contexts.

Two complementary analyses were performed with the fitted models. First, we assessed whether placement errors differed across classifications, computing estimated marginal means per context (*emmeans* package) and comparing contexts pairwise with model-based t-tests (*contrast* function); classifications including the time-of-day smooth (Supplementary Table S3) were excluded to limit comparisons. Second, we tested each context’s mean placement error against zero (no systematic difference between body-worn and glasses-level measurements) using model-based t-tests (*test* function). For interpretability, estimates (log_10_ scale) were back-transformed to percentage errors. Both analyses used α = 0.05.

### Outcome-level analysis

To quantify how placement affects daily summary metrics, each of the 54 metrics was modelled separately with linear or generalised linear mixed-effects models, the engine and family chosen from each response’s distribution^11^. Most metrics (46 of 54) used linear mixed-effects models (Gaussian). Continuous, right-skewed, zero-inflated duration metrics (e.g., total duration above threshold; 5 metrics) used a Tweedie distribution^55^ with a log link, and count metrics (pulses above threshold, frequency of threshold crossings; 3 metrics) a Poisson family. Some responses were transformed before modelling: metrics pre-transformed with the zero-inflated log_10_ transformation were modelled on that scale^51^, interdaily stability was logit-transformed to respect its [0, 1] range, and pulse-level metrics (mean level and duration) used the zero-inflated log_10_ scale^51^. Full specifications are in Supplementary Table S2.

All models shared the same fixed- and random-effects structure, expressed in Wilkinson notation as:

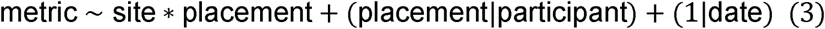

The fixed effects are the measurement *site* (location of data collection), the *placement*, and their interaction, which allows the placement effect to vary across sites. The random-effects structure includes random intercepts and random slopes for *placement* per *participant*, capturing between-participant variability in placement effects, and random intercepts per calendar *date* (participant-day) for day-to-day variability. Glasses was the reference level for placement, so estimates for chest and wrist represent deviations from glasses-level measurements. Site was sum-coded (deviation coding), so the intercept represents the grand mean across sites rather than a single reference.

When models failed to converge or were singular with the full random-effects structure, complexity was reduced stepwise: first the date random intercept was removed, then, if needed, the placement random slope was simplified to an intercept only. All 54 models converged and were non-singular. Diagnostics (*performance* package) checked normality of residuals and random effects, homogeneity of variance, linearity, and collinearity.

Fixed-effect significance was assessed with Wald *χ*^2^ tests (*Anova*, *car* package; α = 0.05). To express placement effects as scale-independent relative differences, estimates were computed relative to the baseline using Equation 4:

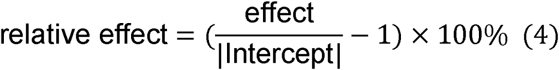

where the intercept is the estimated marginal mean for glasses averaged across sites, and the effect is the estimated difference between a body-worn placement (chest or wrist) and glasses.

Marginal and conditional *R*^2^ were calculated following Nakagawa and colleagues^56^. Partial *R*^2^ contributions were computed as the difference in *R*^2^ between the full model and a reduced model omitting the component of interest, for the placement fixed effect, the site fixed effect, the placement random slope across participants, the date random effect, and the participant random intercept.

Across all 52 metrics with participant-day estimates, grouped into six metric classes (duration, exposure history, level, spectrum, temporal dynamics, and timing), we quantified placement bias as the participant-day difference between a comparison position (chest or wrist) and the glasses reference on each metric’s model scale. Interdaily stability and intradaily variability were excluded because they have no participant-day estimates. We evaluated sampling precision using hierarchical bootstrap resampling (B = 10,000 replicates per design scenario). For the participant axis, 1–200 participants were sampled with replacement, with either 3 or 7 days sampled per selected participant. For the monitoring-duration axis, either 25 or 50 participants were sampled with replacement, with 1–7 days sampled per participant. At each sample size, we calculated the standard deviation of the bootstrap distribution of the mean bias and expressed it as a percentage of the absolute mean model-scale glasses reference for the corresponding metric. Metric-specific curves were retained, while the pointwise median across metrics within each class provided the class-level summary. The crossing analysis reported the first sample size at which this class-median standard deviation was at or below 5%.

## Results

### Measurement-level analysis

Estimated placement errors for each contextual classification are shown in Figure 2. Errors were consistently larger for wrist-than chest-worn dosimeters and generally larger at night than during the day. Across all categorical classifications (environment, activity, and indoor-work lighting) and for nearly all hours, mean errors were negative, indicating that body-worn dosimeters underestimated momentary glasses-level exposure. Almost all errors differed significantly from zero; the exceptions were chest-worn measurements in vehicles during the day (Figure 2, Panel A) and outdoor measurements during the night and morning (Figure 2, Panel B, approximately 02:00–12:00 local time).

**Figure 2:**
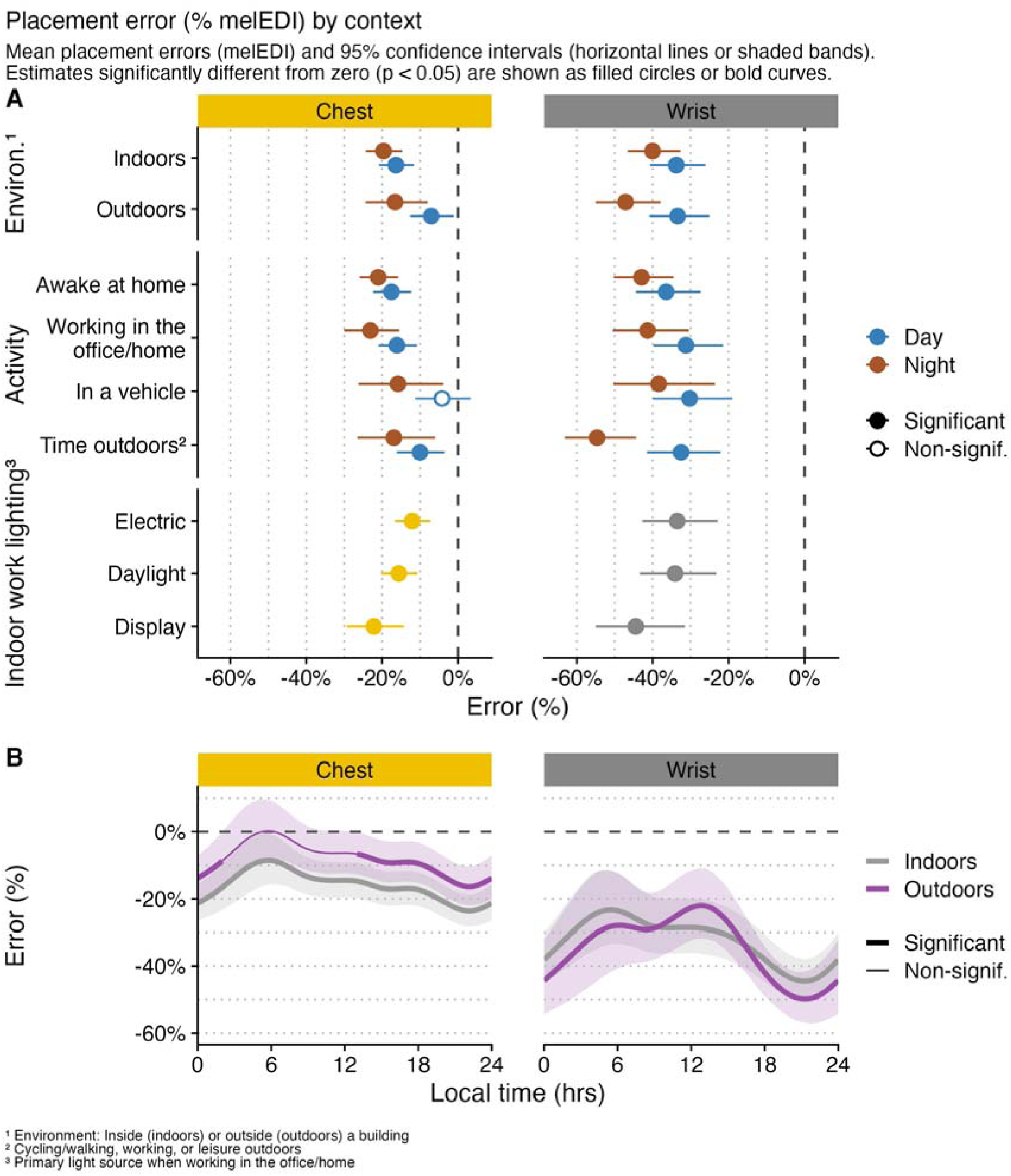
Placement errors (% melEDI) by context. Significant deviations from zero (p < 0.05, unadjusted for multiple comparisons) are marked by filled circles or bold curve segments. An error of 0% indicates no difference between the body-worn and glasses-level measurements, while a negative error indicates that the body-worn dosimeter underestimates momentary glasses-level light exposure.

Differences in placement errors between contexts are shown in Figure 3 (significant differences in bold). For chest-worn dosimeters, 6 of 17 comparisons were significant, versus 7 of 17 for wrist-worn dosimeters. Among classifications evaluated separately by photoperiod (Supplementary Table S3), all significant chest differences occurred during daytime, whereas significant wrist differences occurred during both day and night. For the indoor-work light source, placement errors differed significantly between electric and display lighting at both placements.

**Figure 3:**
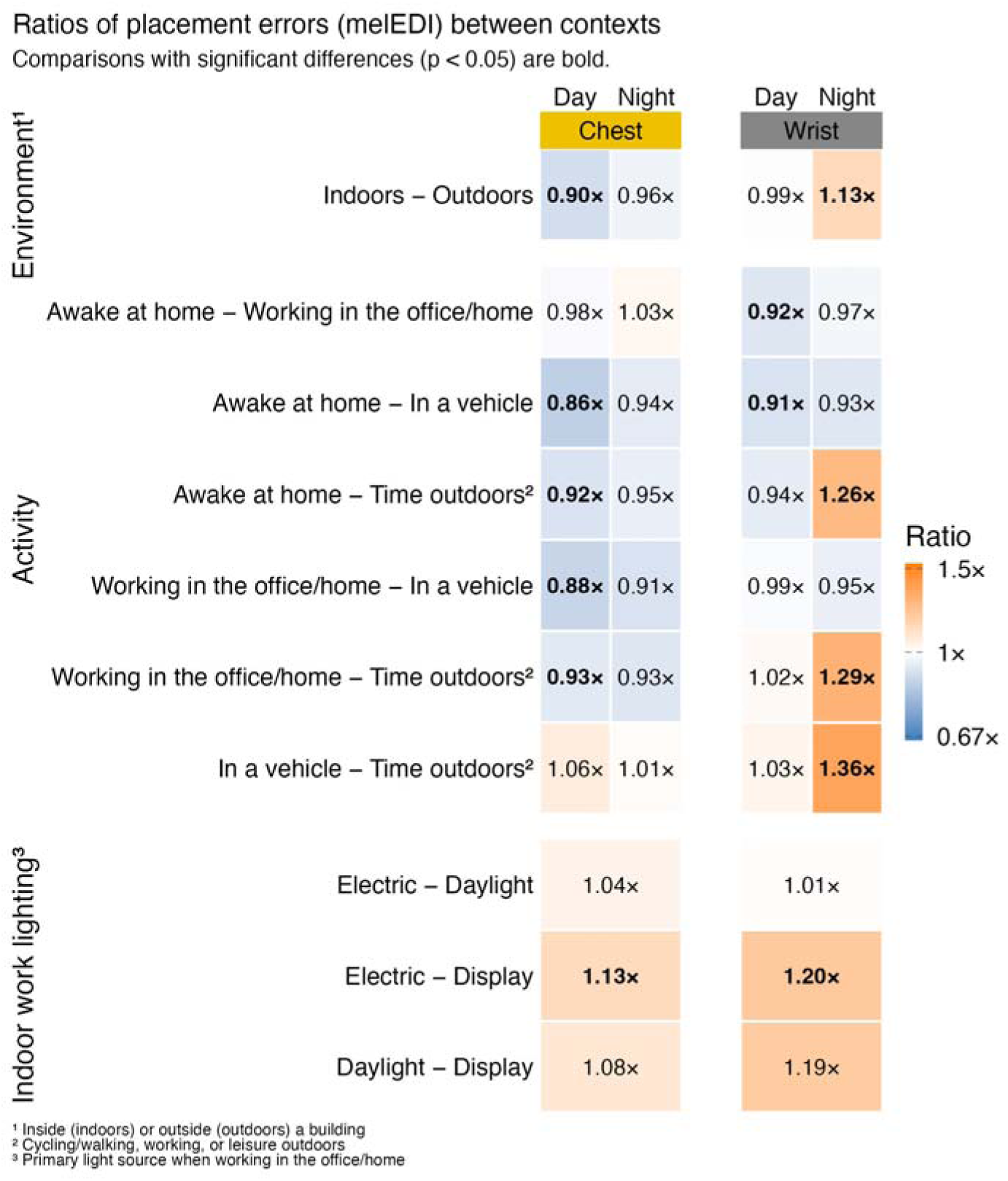
Ratios of placement errors (melEDI) between contexts. Significant differences in placement errors between contexts (p <0.05, unadjusted for multiple comparisons) are shown in bold. The ratio in each cell is the placement error in the second context relative to that in the first context, indicating how many times larger (ratio > 1) or smaller (ratio < 1) the error is in the second context.

### Outcome-level analysis

The effects of placement on the 54 metrics are summarised per metric class in Table 1 and reported in full in Supplementary Table S5. Compared with the glasses-level reference, 19 metrics reached nominal significance at the chest (35%) and 27 at the wrist (50%). Across all metrics, the median absolute bias was 1% at the chest and 2% at the wrist; among nominally significant contrasts, it was 5% at both placements. Thus, the same significant-contrast median applied to a larger share of wrist metrics. The median participant-level standard deviation was approximately 8%. The placement fixed effect accounted for approximately 2% of the variance, versus approximately 26% for participant-level differences. Both placements predominantly underestimated the glasses-level metric: bias was negative for 13 of 19 significant chest contrasts and 16 of 27 significant wrist contrasts.

**Table 1:** Summary overview of outcome-level bias due to dosimeter placement, compared with glasses. Biases are based on each metric’s modelling scale (see Supplementary Table S2). Brightness-based timing metrics are referenced to the participant-weighted mean wake duration (15 h 44 min), darkness-based timing metrics to mean sleep duration (8 h 16 min), and all other metrics to the absolute glasses reference.

| Summary overview of outcome-level bias<br>for body-worn versus eye-level measurement |  |  |  |
| --- | --- | --- | --- |
| Primary analysis: documented non-wear removed; diary-defined sleep retained<br>Timing metrics use mean wake/sleep durations;<br>other metrics to the absolute glasses reference. |  |  |  |
|  | Chest<sup>1</sup> | Wrist<sup>1</sup> | General<sup>1,2</sup> |
| Duration | bias: 1% (0% - 4%)<br>signif: 6/12, direction: +3/-9 | bias: 2% (0% - 8%)<br>signif: 8/12, direction: +2/-10 | R<sup>2</sup><sub>Pos.</sub>: 3% (1%–7%)<br>R<sup>2</sup><sub>Id</sub>: 24% (16%–39%)<br>SD<sub>Id</sub>: ±6% (2–20%) |
| Exposure history | bias: 2% (1% - 2%)<br>signif: 0/2, direction: +1/-1 | bias: 1% (0% - 1%)<br>signif: 0/2, direction: +2/-0 | R<sup>2</sup><sub>Pos.</sub>: ---<br>R<sup>2</sup><sub>Id</sub>: 31% (26%–37%)<br>SD<sub>Id</sub>: ±21% (10–32%) |
| Level | bias: 5% (1% - 50%)<br>signif: 8/9, direction: +2/-7 | bias: 9% (1% - 84%)<br>signif: 7/9, direction: +2/-7 | R<sup>2</sup><sub>Pos.</sub>: 3% (1%–5%)<br>R<sup>2</sup><sub>Id</sub>: 30% (20%–37%)<br>SD<sub>Id</sub>: ±21% (4–137%) |
| Spectrum | bias: 1% (1% - 1%)<br>signif: 0/1, direction: +0/-1 | bias: 5% (5% - 5%)<br>signif: 1/1, direction: +0/-1 | R<sup>2</sup><sub>Pos.</sub>: 2% (2%–2%)<br>R<sup>2</sup><sub>Id</sub>: 35% (35%–35%)<br>SD<sub>Id</sub>: ±11% (11–11%) |
| Temporal dynamics | bias: 12% (0% - 18%)<br>signif: 5/7, direction: +6/-1 | bias: 15% (5% - 52%)<br>signif: 7/7, direction: +6/-1 | R<sup>2</sup><sub>Pos.</sub>: 2% (0%–15%)<br>R<sup>2</sup><sub>Id</sub>: 53% (18%–69%)<br>SD<sub>Id</sub>: ±33% (11–51%) |
| Timing | bias: 1% (0% - 1%)<br>signif: 0/23, direction: +19/-4 | bias: 1% (0% - 3%)<br>signif: 4/23, direction: +21/-2 | R<sup>2</sup><sub>Pos.</sub>: 1% (1%–1%)<br>R<sup>2</sup><sub>Id</sub>: 19% (11%–41%)<br>SD<sub>Id</sub>: ±6% (3–12%) |
| Overall | bias: 1% (0% - 50%)<br>signif: 19/54, direction: +31/-23 | bias: 2% (0% - 84%)<br>signif: 27/54, direction: +33/-21 | R<sup>2</sup><sub>Pos.</sub>: 2% (0%–15%)<br>R<sup>2</sup><sub>Id</sub>: 26% (11%–69%)<br>SD<sub>Id</sub>: ±8% (2–137%) |
| Overall (significant) | bias: 5% (1% - 50%)<br>direction: +6/-13 | bias: 5% (1% - 84%)<br>direction: +11/-16 | R<sup>2</sup><sub>Pos.</sub>: 2% (0%–15%)<br>R<sup>2</sup><sub>Id</sub>: 26% (11%–69%)<br>SD<sub>Id</sub>: ±8% (2–137%) |

median bias (min - max), significant bias / all metrics, positive / negative deviations
<sup>1</sup> Timing effects and timing-model SDs use the grand mean diary-defined wake duration (15 h 44 min; 1% = 9.4 min) for brightness-based timing metrics and the grand mean diary-defined sleep duration (8 h 16 min; 1% = 5.0 min) for darkness-based timing metrics. All other metrics use the absolute glasses reference on the modelling scale. Overall rows combine these normalization bases.
<sup>2</sup> R<sup>2</sup><sub>Pos.</sub>: partial R<sup>2</sup> for the specific position effect (glasses, chest, or wrist; median min-max), significant metrics only. R<sup>2</sup><sub>Id</sub> and SD<sub>Id</sub>: individual ID random effect, all metrics.

The magnitude of bias varied across metric categories. Among the 12 duration-based metrics, the median absolute bias was 1% at the chest (six significant contrasts) and 2% at the wrist (eight). Neither exposure-history metric differed significantly at either placement. The nine level-based metrics showed more pronounced biases: 5% at the chest (eight significant) and 9% at the wrist (seven). The single spectrum-based metric did not differ significantly at the chest but showed a significant 5% bias at the wrist. The largest median biases were for the seven temporal-dynamics metrics: 12% at the chest (five significant) and 15% at the wrist (all seven significant). By contrast, the 23 timing-based metrics were comparatively insensitive to placement. After normalising brightness-based timing metrics to the participant-weighted mean wake duration (15 h 44 min; 1% = 9.4 min) and darkness-based timing metrics to the corresponding mean sleep duration (8 h 16 min; 1% = 5.0 min), the median absolute timing bias was 1% at both placements, with rounded ranges of 0%–1% at the chest and 0%–3% at the wrist. No timing metric differed significantly at the chest, whereas four differed at the wrist.

Duration- and level-based metrics tended towards underestimation, with most contrasts negative, whereas temporal-dynamics metrics tended towards overestimation, with most contrasts positive.

Among metrics with significant placement effects, absolute biases ranged from 1% to 50% at the chest and 1% to 84% at the wrist, both maxima arising for the *Dark threshold* of the *Barroso metrics*^35^. Distributions of signed and absolute biases for significant and non-significant contrasts are shown in Figure 4.

**Figure 4:**
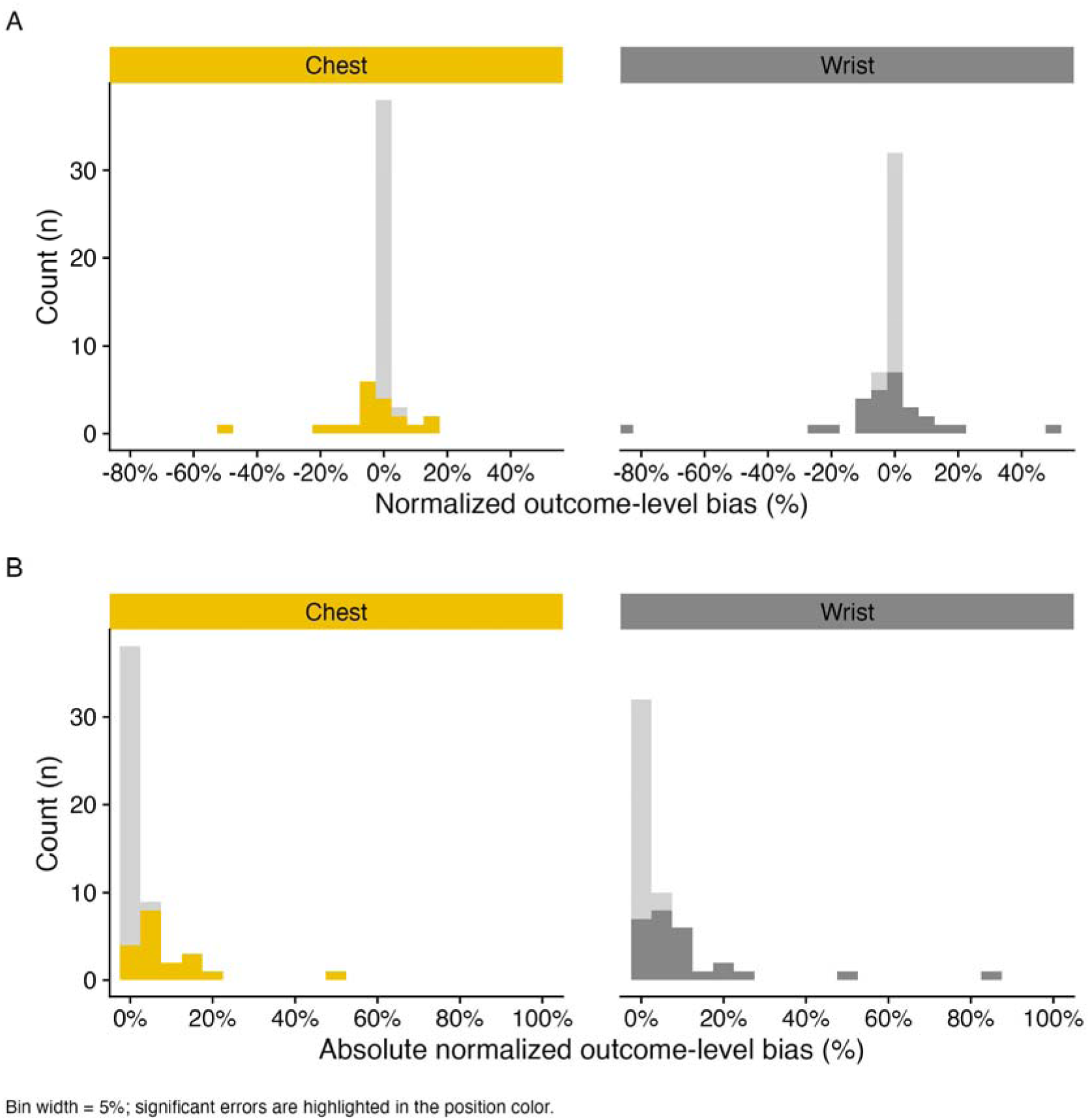
Distribution of the (A) outcome-level bias across metrics and (B) absolute bias. Significantly deviating metrics are shown in the position colour, non-significant deviations in light grey.

Site-by-placement interactions were detected for 12 of the 54 metrics, indicating that placement bias was not fully consistent across sites (Supplementary Table S5). Of the 432 site–metric combinations (54 metrics × 8 sites), chest biases exceeded the across-site average in 14 instances and were smaller in 11; wrist biases exceeded it in 16 and were smaller in 10. In total, site effects mediated the placement effect in 51 of 432 cases (∼12%).

The random effect of date could be estimated for 11 of the 54 metrics. For these, day-to-day variability was substantial (date-level standard deviations approximately 6%–65%), and the partial variance contribution of date consistently exceeded that of the placement fixed effect: partial *R*^2^ 13%–27% for date versus 0%–5% for placement.

Participant-specific random slopes for placement could be estimated for 12 of 54 metrics. Participant-specific variation in the placement effect was considerable, though smaller than the date effect, with random-slope standard deviations ranging from less than 1% to 59% at the chest and from 1% to 80% at the wrist. In all but one such model, the random-slope standard deviation exceeded the corresponding population-average placement bias. Thus, although the average placement bias was generally modest, its magnitude varied substantially among participants and is expected to switch sign (e.g., from under- to overestimation) with some regularity: assuming the random-slope standard deviation equals the fixed-effect bias and disregarding random intercepts, about 16% of participants would have a bias opposite to the average. Population-average correction factors would therefore not necessarily correct individual participants accurately.

As an exploratory sensitivity analysis, the pipeline was repeated without excluding documented non-wear periods, giving a tentative indication of how the treatment of non-wear affected estimates. Results were broadly consistent with the primary analysis (Supplementary Table S6), though estimated placement biases were generally smaller when non-wear periods were retained. This attenuation should not be read as improved agreement with the glasses-level reference, because retaining these periods introduces an additional source of measurement bias from device non-wear.

The wake-only sensitivity analysis, which excluded documented sleep in addition to non-wear, produced results broadly consistent with the primary analysis (Supplementary Table S7). Across the reduced set of 42 metrics available in both analyses, the median absolute dosimeter-placement bias remained 1% at both the chest and wrist, while the number of significant contrasts changed only slightly, from 12 to 11 at the chest and from 16 to 15 at the wrist. The upper tail of the bias distribution was attenuated, however: the maximum absolute bias decreased from 15% to 10% at the chest and from 25% to 16% at the wrist, with the largest reduction observed for mean melanopic EDI. Thus, the overall placement pattern was robust to the removal of sleep observations, although sleep-time measurements appear to contribute to some of the largest deviations; this comparison is conditional on the waking-wear metric set because nighttime-focused metrics were excluded.

Across the 52 metrics with participant-day estimates, the class-median bias SD fell steeply as the number of participants increased, consistent with the expected 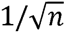 sampling-error curve (Supplementary Figure S1, Supplementary Table S8). With three days per participant, the duration, spectrum, and timing classes reached the 5% tolerance with 3–5 participants, whereas exposure history and level required 7– 21 participants. Temporal dynamics was the least stable class: its median crossed 5% at 45 participants for the chest and 76 for the wrist. Increasing monitoring to seven days reduced these requirements to 33 and 59 participants, respectively, while the other class medians crossed between 2 and 16 participants. Adding days per participant produced a smaller benefit. With 25 participants, all class medians except temporal dynamics reached 5% within one or two days, whereas temporal dynamics did not cross within seven days. With 50 participants, all other class medians were already below 5% after one day; temporal dynamics crossed after three days for the chest but remained above 5% for the wrist after seven days. Individual metrics nevertheless remained heterogeneous, so a class-median crossing does not imply that every metric within that class met the tolerance.

## Discussion

This study examined placement bias at two complementary analytical scales. The measurement-level analysis quantified differences between simultaneous body-worn and eye-level measurements before aggregation, capturing how closely a body-worn dosimeter reproduces momentary eye-level exposure. The outcome-level analysis compared daily summary metrics derived separately at each placement, evaluating the consequences of placement for the exposure variables used in epidemiological or intervention studies, but allowing overestimation during some periods to offset underestimation during others. The two analyses thus address distinct questions: time-resolved proxy error versus the resulting bias in derived outcomes.

### Measurement-level analysis

Body-worn dosimeters underestimated eye-level exposure in almost all contexts: estimated mean placement errors were negative and almost all differed significantly from zero (Figure 2), the exceptions being chest-worn measurements in vehicles during the day and outdoor measurements during the late night and morning (approximately 02:00–12:00 local time). Errors were consistently larger for wrist-than chest-worn dosimeters, ranging across categorical classifications from −23.1% to −4.2% (chest) and −54.7% to −30.3% (wrist). Time-of-day errors spanned a similar but slightly smaller range (−23.6% to 0.2% chest; −49.8% to −22.0% wrist). This likely reflects that the categorical classifications were designed to separate illumination fields and postures, whereas time-of-day errors cut across contexts and therefore average over the environments and activities occurring at a given hour.

Placement errors varied with context, though modestly in absolute terms, spanning just 18.9% (chest) and 24.4% (wrist) across categorical classifications. This is consistent with the pairwise comparisons (Figure 3), where fewer than half were significant, and with the time-of-day errors (Figure 2), where indoor and outdoor confidence intervals overlap. Nonetheless, chest errors reached as little as −4.2% in some contexts, so context could determine whether the error was minor or approached roughly −20%. Wrist errors were at least −30.3% in every context, so context shifted the error but underestimation was already substantial throughout.

Placement errors were consistently larger at night than during the day. A plausible explanation is that, without daylight, illumination comes primarily from electric sources that are more directional and unevenly distributed, producing spatially heterogeneous illumination fields associated with larger placement errors^29,33^. Consistently, among indoor-work light sources the largest errors occurred under display lighting, as displays are fairly directional and expected to produce more heterogeneous illumination fields than electric room lighting or daylight.

For contexts classified by environment and activity, no consistent pattern emerged across placements. For the chest, indoor and outdoor errors differed significantly (Figure 3) during the day (−16.3% indoors, −7.1% outdoors) but not at night (−19.6% vs −16.6%). For the wrist the opposite held: no significant day difference (−33.8% vs −33.4%) but a significant night difference (−40.0% vs −47.1%). Activity-based errors broadly followed the indoor/outdoor pattern, with indoor activities (awake at home, working) resembling the indoor error and outdoor activities (in a vehicle, time outdoors) the outdoor error.

### Outcome-level analysis

Aggregation changed the practical meaning of placement error. Large, predominantly negative momentary errors often averaged to much smaller daily biases, but aggregation did not make the placements interchangeable. Across all 54 metrics, median absolute bias was 1% at the chest and 2% at the wrist. Among nominally significant contrasts, median absolute bias was 5% at both placements, but such contrasts occurred for 19 of 54 metrics (35%) at the chest and 27 of 54 metrics (50%) at the wrist, and the largest biases reached 50% and 84%, respectively. Thus, the identical median among significant contrasts masks both a broader and a more extreme effect at the wrist. Placement should therefore be judged by both the magnitude of bias and the range of outcomes susceptible to it, especially when studies plan secondary analyses beyond a single prespecified metric.

The ordering across metric classes is mechanistically coherent. Timing outcomes were comparatively robust, with a median absolute bias of 1% at both body-worn placements; none of the 23 chest contrasts and four of the 23 wrist contrasts were nominally significant. Under the state-specific normalization, 1% corresponded to 9.4 minutes for brightness-based timing metrics and 5.0 minutes for darkness-based metrics, so the residual shifts were generally on the scale of minutes. By contrast, median absolute bias was 5% at the chest and 9% at the wrist for level metrics, and 12% and 15%, respectively, for temporal-dynamics metrics; five of seven temporal-dynamics metrics were affected at the chest and all seven at the wrist. Duration outcomes were intermediate (1% and 2%), while neither of the two exposure-history metrics was affected. The single spectrum-based metric was not affected at the chest but showed a nominally significant 5% bias at the wrist. Although one metric is insufficient to characterise the sensitivity of the class more broadly, there are few spectrum-based metrics in common use. This ordering is plausible because level metrics inherit attenuation directly, whereas temporal-dynamics metrics can also change when placement alters amplitude, fragmentation, or frequency structure. A body-worn time series may consequently provide a reasonable timing summary while remaining a poor proxy for momentary eye-level exposure or for other outcomes derived from the same measurements.

The state-handling sensitivity analyses support the robustness of this ordering rather than equivalence between placements. Retaining documented non-wear produced apparently smaller biases by mixing valid measurements with periods when devices were not at their intended positions; this is attenuation through additional misclassification, not improved validity. Among the 42 metrics shared with the wake-only analysis, removing sleep left the median absolute bias at 1% for both placements and changed the number of nominally significant contrasts only from 12 to 11 at the chest and from 16 to 15 at the wrist. It nevertheless reduced the maximum absolute bias from 15% to 10% at the chest and from 25% to 16% at the wrist, with the largest reduction for mean melanopic EDI. This suggests that bedside measurements during recorded sleep contributed to some extreme daily differences without driving the broader placement pattern. Because nighttime-focused metrics were unavailable in the wake-only analysis, this reassurance applies only to outcomes that can be defined from valid waking wear.

Variation across sites, days, and participants is more consequential for translation than the modest average bias of many metrics. Across outcomes, the placement fixed effect accounted for approximately 2% of the variance, compared with approximately 26% for participant-level differences; site mediated the placement effect in 51 of 432 site–metric cases (about 12%). Date-level standard deviations ranged from 6% to 65%, while participant-specific placement-effect standard deviations ranged from less than 1% to 59% at the chest and from 1% to 80% at the wrist, exceeding the corresponding population-average placement effect in all but one case. Participant-specific effects could also differ in direction from that average. A population correction may therefore improve a group mean without recovering eye-level exposure for an individual. Increasing the sample size improves precision in the population-average effect; it does not remove individual measurement error.

The bootstrap converts this heterogeneity into study-design guidance (Supplementary Figure S1, Supplementary Table S8). With seven monitoring days and 10,000 bootstrap draws, the median metric in each non-temporal-dynamics class reached the 5% precision criterion with 2–16 participants, whereas temporal-dynamics outcomes required 33 participants at the chest and 59 at the wrist. Increasing participant numbers was generally more effective than adding monitoring days because longer monitoring reduces within-participant sampling uncertainty but cannot average away between-participant differences. The 5% criterion describes uncertainty in the estimated mean bias—not an acceptable magnitude of bias—and the class medians conceal substantial metric-level variation. These empirical projections should therefore inform validation-subsample design around the intended metric, rather than be treated as universal sample-size requirements.

### Literature context

Our results can be compared with previous studies that assessed placement bias with similar methods, placing dosimeters near the eyes and on the body and comparing the measurements.

Few studies report momentary (measurement-level) differences, as most aggregate over time. Aarts et al.^26^ reported smaller median deviations at the chest than the wrist and slightly larger errors indoors than outdoors, broadly consistent with our results. Wen et al.^27^, by contrast, found eye-level measurements significantly lower than chest and wrist (a positive bias). Because of the high variance of the placement error, studies with few participants may reach different conclusions from ours (Supplementary Figure S1). More recently, de Vries et al.^29,33^ quantified placement errors of chest-worn dosimeters under simulated indoor-work lighting, reporting a range of errors depending on the illumination field and posture. Under idealised overhead electric lighting, chest dosimeters overestimated eye-level measurements, whereas underestimation occurred under vertically oriented sources such as windows or displays. Their results for vertical sources (Figure 3 in^33^) broadly agree with ours for daylight and display lighting (Figure 2). However, whereas most of their ceiling-only simulations indicated overestimation, we observed chest underestimation when electric light was the primary source, though smaller than under daylight or display lighting. This likely reflects that real-world electric lighting is not strictly ceiling-based and that our light-source classification is only indicative, as secondary sources may also have been present.

Direct comparison of our outcome-level findings with the existing literature is limited because previous studies evaluated different optical quantities, summary metrics, thresholds, time periods, and sensor systems (Supplementary Table S1). Nevertheless, the available studies support the conclusion that aggregation into daily metrics can reduce, but does not eliminate, placement-dependent differences. Bhandari et al.^25^ reported wrist values approximately 37–38% lower than temple-level measurements for mean daily illuminance, but only about 22% lower for time above 1,000 lx. This contrast is consistent with our finding that level-based metrics were generally more placement-sensitive than duration- or timing-based outcomes. Similarly, Cole et al.^23^ observed almost identical 24-hour mean illuminance at the forehead and wrist despite identifiable discrepancies in the underlying time series, illustrating how positive and negative deviations may cancel during daily aggregation. In contrast, Wen et al.^27^ found both mean illuminance and the proportion of time above 1,000 lx to be approximately twice as high at the chest and wrist as at eye level, demonstrating that aggregation does not ensure small bias, when placement differences are systematic within the monitored setting. Figueiro et al.^24^ likewise reported substantial placement dependence in absolute photopic and circadian exposure despite greater similarity in broad temporal patterns. Taken together, these studies broadly agree with our observation that the validity of a body-worn proxy is metric-specific: outcomes determined mainly by exposure level or amplitude are generally more vulnerable to placement bias, whereas some timing and duration measures may remain comparatively stable. However, the direction and magnitude of published effects vary considerably, and several confounded placement with dosimeter type or were restricted to a single setting or participant. Our results extend this literature by comparing multiple placements, using one dosimeter model, across a larger, geographically diverse sample and by showing systematically, across 54 metrics, that agreement for one exposure summary cannot be generalised to other metrics derived from the same time series.

A modest systematic underestimation of eye-level exposure is not necessarily undesirable. Illuminance or melanopic EDI measured at the corneal plane can overestimate the effective light input to the eye, because planar measurements do not account for the restricted, directionally non-uniform human field of view^40,57,58^. In principle, a small negative placement bias could partially offset this geometric overestimation. However, the present results do not support treating chest- or wrist-level underestimation as a reliable compensatory correction: the bias varied substantially between participants, metrics, contexts, and days, and individual effects could differ in size and direction from the average. Any compensation would therefore be incidental, and a population-level correction could reduce error for some individuals while increasing it for others.

### Limitations

Several limitations relate to the study population, protocol and interpretation of the eye-level reference. Participants were predominantly healthy adults whose daily routines commonly involved indoor work. Placement effects may differ in children, older adults, clinical populations, outdoor workers or groups with systematically different postures, mobility patterns, clothing or occupational environments. Sample sizes also varied between sites and were comparatively small at UCR and TUM, with six and ten participants, respectively. Although the multisite design increased the diversity of environments represented, estimates of site-specific deviations are therefore less precise for these locations.

The spectacle-mounted dosimeter provided a practical glasses-level reference but did not measure corneal or retinal exposure directly. Its position, orientation, and field of view differ from those of the eye, and it cannot account for eye rotations, eyelid closure, pupil size, or transmission through the ocular media. The glasses-level reference should therefore be interpreted as an approximation of light incident near the corneal plane rather than a direct measurement of retinal irradiance.

Nevertheless, because the three dosimeters were of the same type and recorded simultaneously, the comparisons isolate anatomical placement more directly than studies confounding device type with placement.

Non-wear periods were identified from participant records and could be removed only when documented. Undocumented removal, displacement, or occlusion may remain in the dataset and contribute to both measurement- and outcome-level differences. The sensitivity analysis retaining documented non-wear periods produced generally smaller placement biases, illustrating that apparent agreement can improve when a separate source of misclassification is introduced; this should not be read as evidence that retaining non-wear improves validity.

For daily metrics, recorded sleep periods were retained even though the spectacle-mounted and chest-worn dosimeters were not worn during sleep. Participants placed these devices face-up on a bedside surface, so they measured ambient light in the sleep environment rather than light at their habitual placements; the wrist dosimeter may also have been covered by bedding or clothing. Metrics incorporating sleep therefore mix placement-specific measurements during wakefulness with differently situated environmental measurements during sleep, most relevantly for metrics focused on nocturnal light levels.

The contextual classifications introduce additional uncertainty, as each combines a heterogeneous range of activities, postures, and illumination environments; placement errors in specific situations may therefore exceed the category-average estimates, because within-category variation is smoothed. Contextual information came from hourly logbooks that have been completed retrospectively, and activity or illumination transitions may not align with hourly boundaries. Misclassification and category overlap would generally attenuate differences between contexts, so the analysis may underestimate how strongly specific postures or illumination configurations modify placement errors.

As noted in the Methods, modelling placement error is challenging because the response is the difference between two paired, noisy measurements and is strongly heavy-tailed. The GAMMs were specified to accommodate this, but residual diagnostics indicated that the tails remained incompletely represented; the estimated mean placement errors underlying our conclusions are likely less sensitive to this misspecification than tail probabilities, but confidence intervals and hypothesis tests should be regarded as approximate. The analyses also included many related contrasts and metrics without correction for multiple comparisons, and were therefore treated as estimation-focused and exploratory rather than independent confirmatory tests. Some nominally significant findings may be false positives, and interpretation should emphasise effect magnitude, uncertainty, directional consistency, and replication across related metrics or contexts rather than individual p-values.

Finally, fewer observations were available at night than during the day (see Supplementary Figure S2), because sleep-time measurements were excluded from the measurement-level analysis. Nighttime estimates are therefore less precise and may be disproportionately influenced by the remaining waking observations; they should not be read as representative of the full nocturnal period, which is dominated by sleep.

### Practical implications

The choice of placement should be guided by the primary outcome, expected duration, sample size, and the practical risk of occlusion or non-wear. Eye-level placement remains preferable when momentary exposure values are the principal outcome, when temporal-dynamics metrics are of primary interest, for individual-level inference in which participant-specific error cannot be averaged away, or for small group samples in which the population-average placement effect remains imprecise. It is also important in short protocols, where the added burden of spectacle-mounted instrumentation is limited, and in environments where directional or heterogeneous illumination is expected to produce large differences between placements. However, spectacle-mounted measurements have their own limitations, including occlusion, displacement, and reduced acceptability during extended monitoring.

Chest-worn dosimeters offer the most defensible compromise when eye-level measurement is not feasible. Across all metrics, typical outcome-level bias was smaller and the upper tail less extreme at the chest than at the wrist, although the median among nominally significant contrasts was 5% at both placements and fewer chest metrics were affected. Chest placement may also support better adherence during long-term monitoring^14,18^, making it suitable for extended campaigns and larger samples where burden, device retention, and scalability outweigh the need to reproduce momentary eye-level exposure. It may be comparatively defensible when duration, exposure-history, or timing outcomes—or the single spectrum outcome examined—are primary, because these were less placement-sensitive than level or temporal dynamics. Nevertheless, “chest-worn” spans a range of locations and orientations; bias may be reduced by orienting the dosimeter so its measurement plane approximates the habitual viewing direction^29,33^, which may also reduce between-participant heterogeneity in similar contexts^32^.

From a measurement-error perspective, the findings give little reason to prefer wrist placement when eye-level or chest placement is viable: wrist measurements showed larger and more consistent momentary underestimation, nominal differences across a larger proportion of metrics, more extreme outcome-level estimates, and lower precision for population-average temporal-dynamics bias under the evaluated designs. Wrist placement may remain defensible when burden or adherence requires it and timing outcomes are narrowly prespecified, but four wrist timing metrics still differed nominally, and relatively good timing performance does not validate level, duration, temporal dynamics, or the underlying time series. Wrist placement should therefore not be selected solely for convenience when eye-level or chest placement is feasible and broad secondary, level, or temporal-dynamics analyses are anticipated.

For studies seeking to quantify population-average placement bias, recruiting more participants should generally take priority over extending monitoring. With seven days per participant, the class-median 5% precision tolerance was reached with 2– 16 participants for all classes except temporal dynamics, which required 33 participants at the chest and 59 at the wrist. These are design-specific precision benchmarks, not power requirements or thresholds for acceptable bias. These recommendations assume that documented non-wear is removed and sleep handling is defined prospectively; the wake-only sensitivity analysis supported the pattern for waking-wear outcomes but did not evaluate nighttime-focused metrics.

Measurement context modified placement error, but these contextual effects were generally smaller than the overall difference between placements, especially at the wrist. The implication is not that every analysis must apply detailed contextual correction, but that a limited set of “high-risk” conditions should inform design and interpretation. Figure 2 can be used to approximate the expected direction and magnitude of error for chest- and wrist-worn dosimeters under the activities and environments studied. Larger errors appeared more likely in directional or heterogeneous illumination, including display-dominated settings and some outdoor nighttime conditions, though the illumination field’s spatial properties were not measured directly. Eye-level measurement should be prioritised when such environments are central to the research question, whereas chest placement may be adequate in more homogeneous conditions.

We do not recommend a universal correction factor for chest- or wrist-derived measurements. Although average placement effects were predominantly negative, their magnitude varied across metrics, contexts, days, sites, and participants, and individual effects can differ in direction from the average, so a single multiplicative correction could improve agreement for some observations while worsening it for others. Investigators should instead select placement prospectively according to the intended outcome, report the anatomical location and dosimeter orientation precisely, and interpret findings in light of the metric- and context-specific sensitivity shown here. Where possible, validation subsamples with simultaneous eye-level and body-worn measurements would provide a more defensible basis for quantifying placement error in a given population and setting and should be sized around the least stable prespecified outcome rather than the overall median.

## Acknowledgements

The authors would like to express gratitude to the participants who volunteered in this study, whether they completed the experiment or not.

## Statements

### Ethical Approval

Ethical approval for the overarching study protocol was obtained from the Ethics Committee of the Technical University of Munich (2023-115-S-KK), with local implementation procedures harmonised across participating centres. Additional ethical approvals were obtained from the respective research ethics committees at participating study sites.

### Statement of Author contributions

Conceptualization: JZ, SWV, AD, JD, MS

Methodology: JZ, SWV, AD, JD, MS

Software: JZ, SWV

Data curation: JZ

Formal analysis: JZ, SWV

Investigation: JZ, AD, MS

Resources: AD, JD, MS

Validation: JZ, SWV, AD, JD, MS

Visualization: JZ, SWV

Writing – original draft: JZ, SWV

Writing – review and editing: All authors

Funding acquisition: AD, JD, MS

Project administration: AD, JD, MS

Supervision: JD, MS

All authors reviewed and approved the final manuscript.

### Statement of data availability

All data used in this study are available at https://github.com/MeLiDosProject and are archived on Zenodo (https://zenodo.org/communities/22nrm05_melidos/), accessed through the melidosData R package (https://melidosproject.github.io/melidosData/).

All analyses are available on GitHub (https://github.com/tscnlab/ZaunerDeVriesEtAl_bioRxiv_2026), hosted on GitHub Pages (https://tscnlab.github.io/ZaunerDeVriesEtAl_bioRxiv_2026/), and archived on Zenodo (https://doi.org/10.5281/zenodo.21533915).

The individual site datasets are available from:

| Institution (site<br>Abbr.) | City | Country | Repository | DOI |
| --- | --- | --- | --- | --- |
| KNUST | Kumasi | Ghana | AkuffoEtAl_Datas<br>et_2025 | 10.5281/zenodo.1<br>5576731 |
| UCR | San José | Costa Rica | Sancho-SalasEtAl_Datase<br>t_2025 | 10.5281/zenodo.1<br>7289456 |
| IZTECH | Izmir | Turkey | DidikogluEtAl_Dat<br>aset_2025 | 10.5281/zenodo.1<br>6568109 |
| FUSPCEU | Madrid | Spain | BaezaEtAl_Datas<br>et_2025 | 10.5281/zenodo.1<br>6834951 |
| TUM | Munich | Germany | HildenEtAl_Datas<br>et_2025 | 10.5281/zenodo.1<br>6893901 |
| MPI | Tübingen | Germany | GuidolinEtAl_Dat<br>aset_2025 | 10.5281/zenodo.1<br>6895188 |
| BAUA | Dortmund | Germany | BroszioEtAl_Data<br>set_2025 | 10.5281/zenodo.1<br>8111232 |
| THUAS | Delft | The Netherlands | AertsEtAl_Datase<br>t_2025 | 10.5281/zenodo.1<br>7979893 |
| RISE | Borås | Sweden | NilssonTengelinEt<br>Al_Dataset_2026 | 10.5281/zenodo.1<br>8925834 |

### Statement of AI use

During manuscript preparation, generative artificial intelligence tools (ChatGPT 5.5 and Anthropic Claude Opus 4.8) were used to support language editing, structural refinement, and condensation of text. AI assistance was used to improve clarity, coherence, concision, and journal fit, but **not** to generate, analyse, or interpret primary data. All AI-assisted text was critically reviewed, edited, and approved by the authors, who take full responsibility for the content, accuracy, integrity, and conclusions of the manuscript.

During data analysis, AI assistance was used to bugfix code sections, support figure and table styling, and resolve modelling convergence and singularity issues. AI assistance was **not** used to develop the analysis strategy, write analysis scripts, or interpret results, **except** for the two-directional bootstrap function, which was used in the explorative sensitivity analysis on participant and participant-day numbers. The function was reviewed and approved by JZ.

## Supplementary materials

**Supplementary Figure S1.**
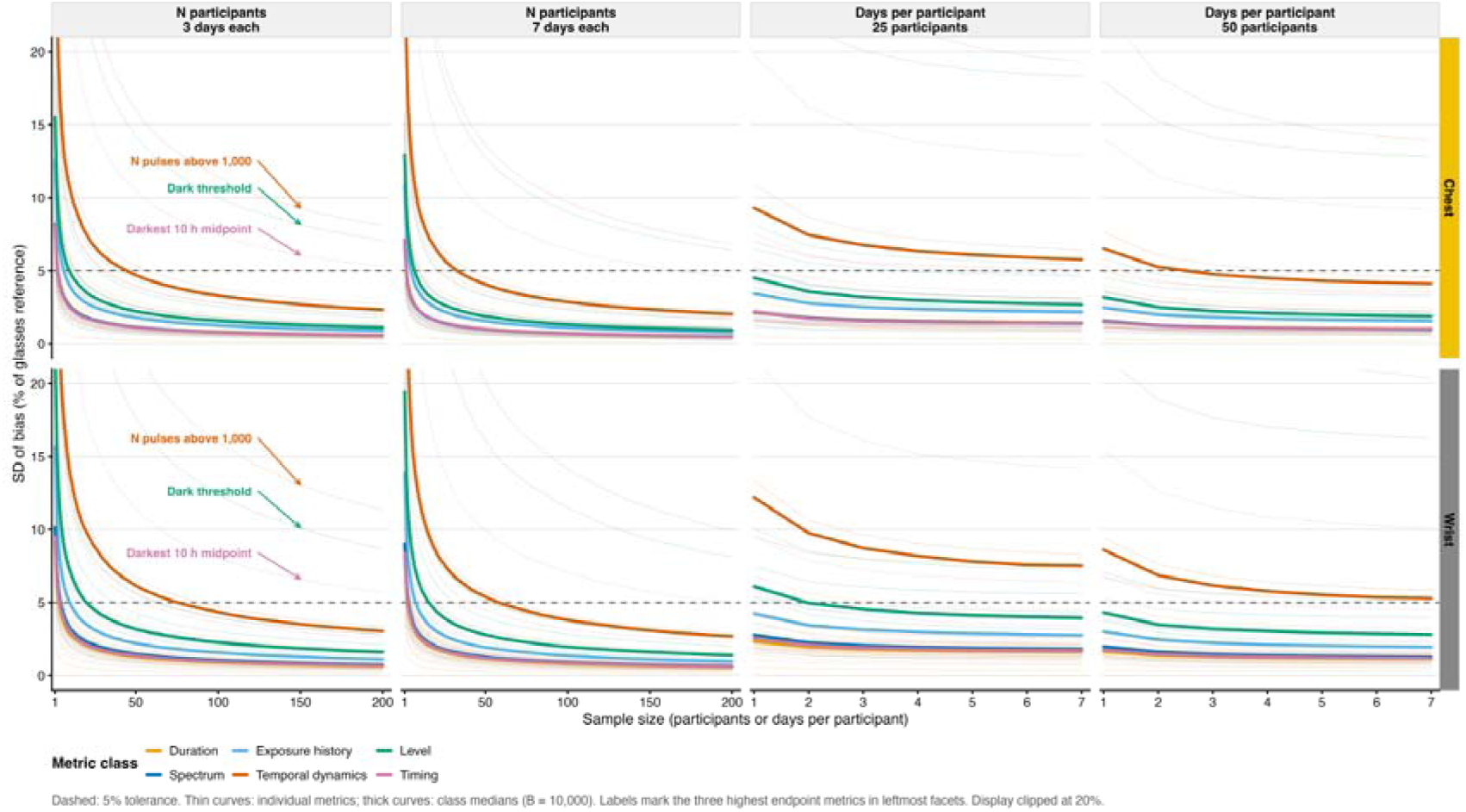

**Supplementary Figure S2.**
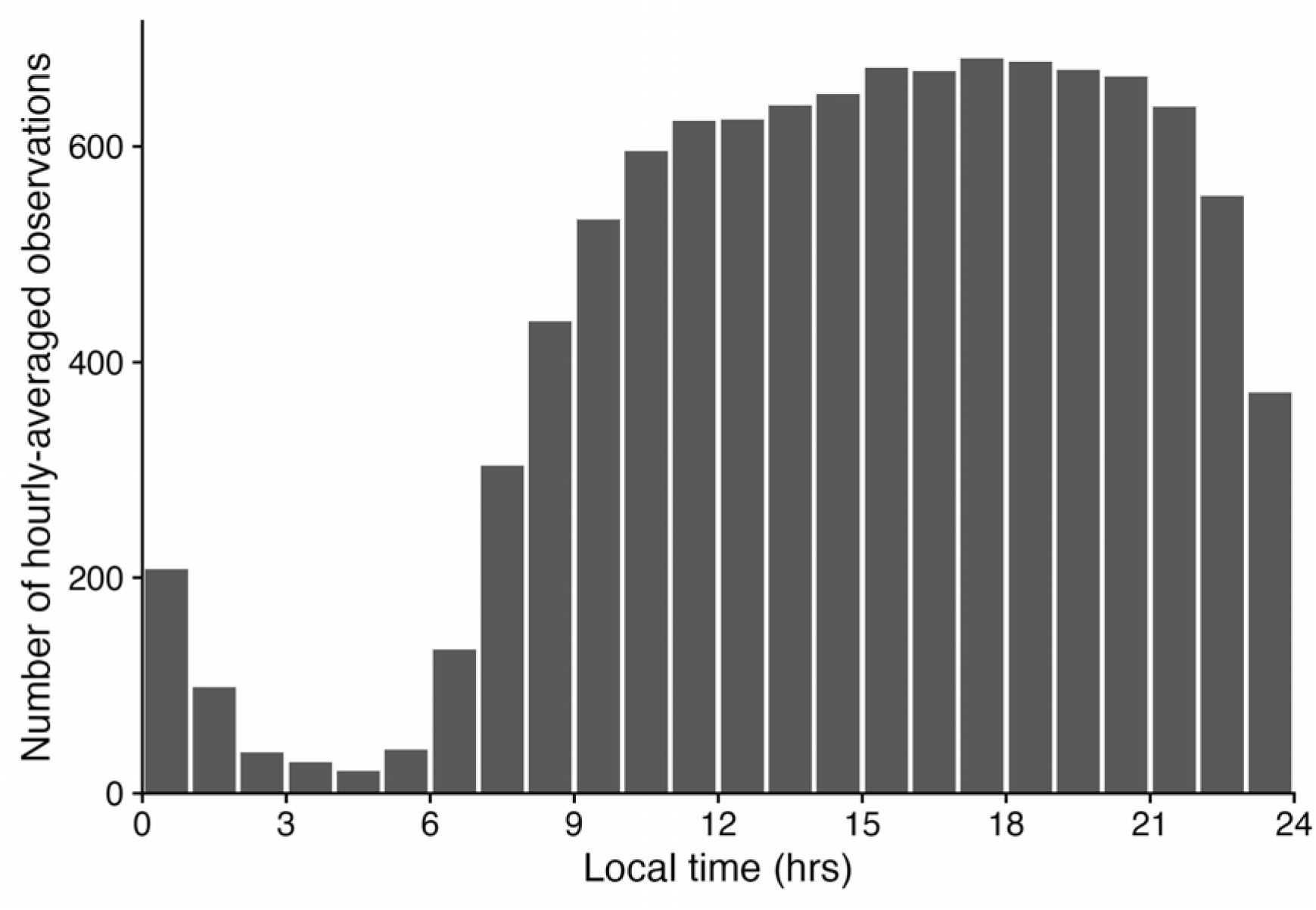

**Supplementary Table S1:** Comparison of placement-dependent differences in personal light and ultraviolet-radiation measurements, focusing on eye- or head-level, chest, and wrist measurements.

| Study | Sample and duration | Measure and outcome | Positions | Position-dependent result | Notes |
| --- | --- | --- | --- | --- | --- |
| Diffey et al. (1977)<sup>19</sup> | One manikin; 2 h each on 19 days | Solar UVR; dose relative to vertex | Vertex; lower sternum | Sternum received 58–73% of vertex exposure, depending on cloud condition. | Rotating unclothed manikin; no behaviour or clothing. No wrist measurement, but hand. |
| Holman et al. (1983), occupational field study<sup>20</sup> | 5 participants; 6 h/day for 11–15 days | Solar UVR; proportion of ambient dose | Vertex; lower sternum; dorsum of hand | Vertex generally received the highest and sternum consistently lower exposure. Hand exposure varied strongly by occupation, from 0.07 of ambient exposure in a classroom teacher to 0.78 in a bricklayer. | One participant per occupation; hand used as the closest approximation to wrist. Sensors were attached to skin, hair, or clothing. |
| Okudaira et al. (1983)<sup>22</sup> | 10 participants; ≥ 24 h | Photopic illuminance | Forward-facing forehead, above the eyes; wrist, oriented like a wristwatch dial | Mean within-person correlation of 0.76 between wrist and forehead log illuminance, with a range of 0.59–0.90 across participants. | Limited reporting of device equivalence and agreement statistics. |
| Cole et al. (1990)<sup>23</sup> | 10 participants; continuous 24 h measurement | Illuminance; minute-level log-lux correlation and 24 h mean illuminance | Forehead-mounted sensor, parallel to forehead; wrist | Wrist and forehead showed closely corresponding temporal patterns: mean within-participant correlation $r = 0.93$ (range 0.88–0.98). Group mean 24 h illuminance was 1,058 lx at the wrist versus 1,061 lx at the forehead, a difference of –3 lx (–0.3%). Participant-level differences or limits of agreement were not reported. | Conference abstract. The forehead position was a proxy rather than a direct eye-level measurement. Devices were randomly interchanged between positions to reduce inter-device bias. Discrepancies occurred when the wrist sensor was covered by bedding after sunrise. |
| Thieden et al. (2000), One-day Beach Study<sup>21</sup> | 11 adults; 5 h | Erythema-weighted UVR; accumulated dose in SED | Top of head; chest; wrist | Mean dose was $19.7 \pm 3.2$ SED at the head, $9.8 \pm 1.3$ SED at the wrist, and 6.6 SED at the chest. Wrist and chest doses therefore averaged $50\% \pm 12\%$ and $34\%$ of head dose, corresponding to mean underestimations of approximately 50% and 66%, respectively. Individual wrist-to-head ratios were approximately 35–76%. | Same dosimeter model and type at all positions. Chest measurements were substantially more variable between participants (CV 54%) than head (16%) or wrist (19%). Over this short period, the wrist-to-head ratio was not constant across individuals, so wrist dose could not precisely estimate head dose. Head placement was on top of a cap and represented maximal cranial rather than ocular exposure. |
| Thieden et al. (2000), Holiday Study<sup>21</sup> | 9 adults; mean 14 days, range 8–26 days | Erythema-weighted UVR; accumulated dose in SED | Top of head; wrist | Mean accumulated dose was $58 \pm 56$ SED at the head and $28 \pm 24$ SED at the wrist. Wrist dose averaged $51\% \pm 15\%$ of head dose, corresponding to a mean underestimation of approximately 49%. Individual wrist-to-head ratios were approximately 34–73%. Wrist and head doses were strongly correlated ($r = 0.89$, $p < 0.01$); the reported regression constrained through zero was head SED = $2.08 \times$ wrist SED. | Same dosimeter model at both positions, although a higher-capacity VioSpor type was used than in the beach study. No chest measurement. The authors interpreted the more stable ratio over the longer period as possible averaging-out of differences in movement and orientation, but duration was not isolated experimentally. Head placement was on top of a cap, not at eye level. |
| Figueiro et al. (2013)<sup>24</sup> | 8 older adults; 5 days | Photopic and circadian light metrics; daily exposure | Near eye; lapel or chest; wrist | Wrist measurements differed substantially from near-eye measurements in absolute exposure, although broad temporal patterns were more similar. | Position and device were partly confounded; devices differed in spectral and directional response. |
| Aarts et al. (2017)<sup>26</sup> | 1 participant; 30 min indoors and 30 min outdoors | Photopic illuminance; deviation from eye-level reference | Between eyes; chest; wrist | Wrist deviation from the between-eyes reference was approximately 27% indoors and 11% outdoors. Chest deviation also depended on environment and activity. | Identical simultaneous sensors, but only one participant and one hour of prescribed activity. |
| Bhandari et al. (2021)<sup>25</sup> | 25 adults; 7 days | Illuminance; daily mean and time $\geq 1000$ lx | Spectacle-mounted at the temple; wrist | Mean daily illuminance was approximately 342–347 lx at the temple and 215 lx at the wrist; the wrist value was therefore about 37–38% lower. The difference was 33% on weekdays (299 vs 200 lx) and 45% on weekends (451 vs 250 lx). Bland–Altman mean difference, Clouclip minus Actiwatch: 126 lx (95% limits of agreement: –160 to 413 lx); daily means were correlated ($R^2 = 0.78$). Time $\geq 1000$ lx was 0.9 versus 0.7 h/day, corresponding to about 12 min/day or 22% less at the wrist ($p = 0.02$); weekday–weekend differences ranged from approximately 13% to 25% lower. | Position and device were fully confounded: Clouclip at the temple versus Actiwatch at the wrist. Devices differed in spectral response, sampling, orientation sensitivity, and measurement range; the wrist sensor could also be shaded by clothing. Differences therefore cannot be attributed solely to sensor position. |
| Wen et al. (2023)<sup>27</sup> | 29 adults; 1 day, mean effective recording time 11.11 ± 3.31 h | Photopic illuminance; mean illuminance and proportion of time >1000 lx | Spectacle-mounted eye level; chest; wrist | Mean illuminance was 189 lx at eye level, compared with 491 lx at the chest and 484 lx at the wrist. Chest and wrist values were therefore approximately 2.59-fold and 2.56-fold higher than eye-level values, corresponding to mean differences of about +302 lx (+160%) and +295 lx (+156%), respectively. Reported pointwise differences ranged from −10,239 to 28,646 lx for eye versus chest and from −10,957 to 32,671 lx for eye versus wrist. The proportion of time >1000 lx was 3.9% at eye level versus 7.8% at the chest and 7.7% at the wrist. Chest and wrist did not differ significantly. | Eye level was measured with a Clouclip, whereas chest and wrist were measured with HOBO sensors, so device and position were partly confounded. In a separate co-location test, Clouclip readings were approximately 1.09 × HOBO + 82.62 lx. Participants followed habitual activities, but monitoring lasted only 1 day and device orientation was actively controlled. |
| Gibaldi et al. (2024)<sup>28</sup> | 1 adult; four 5-min activities (20 min total) | Six-channel spectral irradiance; activity-specific median exposure | Helmet-mounted near eye level; wrist | Wrist irradiance was 124% higher than head-level irradiance on average (2.24-fold) and was more variable. The largest reported difference occurred during smartphone use, when wrist irradiance was 401% higher (5.01-fold). Exact differences for the other activities were not reported numerically. | Identical AS7262 sensors were worn simultaneously, reducing device confounding. Very small exploratory comparison with one adult and brief prescribed activities; the head sensor was close to the eyes but pitched 18° downward and was not an ocular-plane measurement. |

**Supplementary Table S2:**
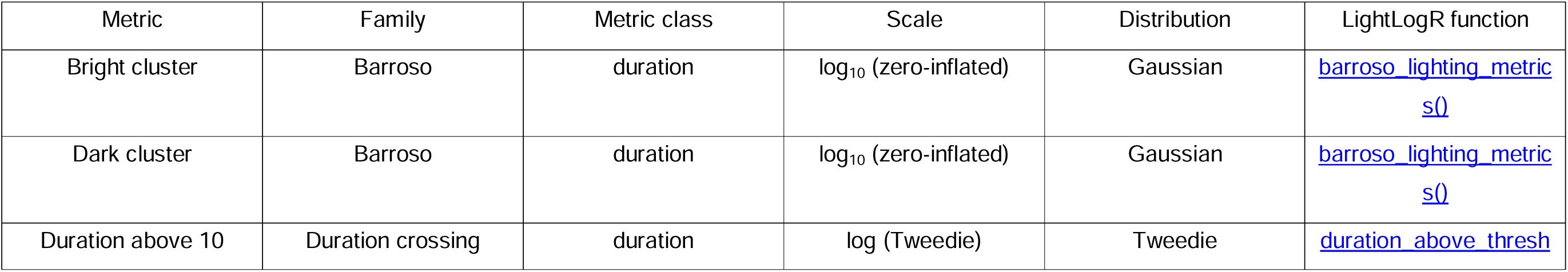
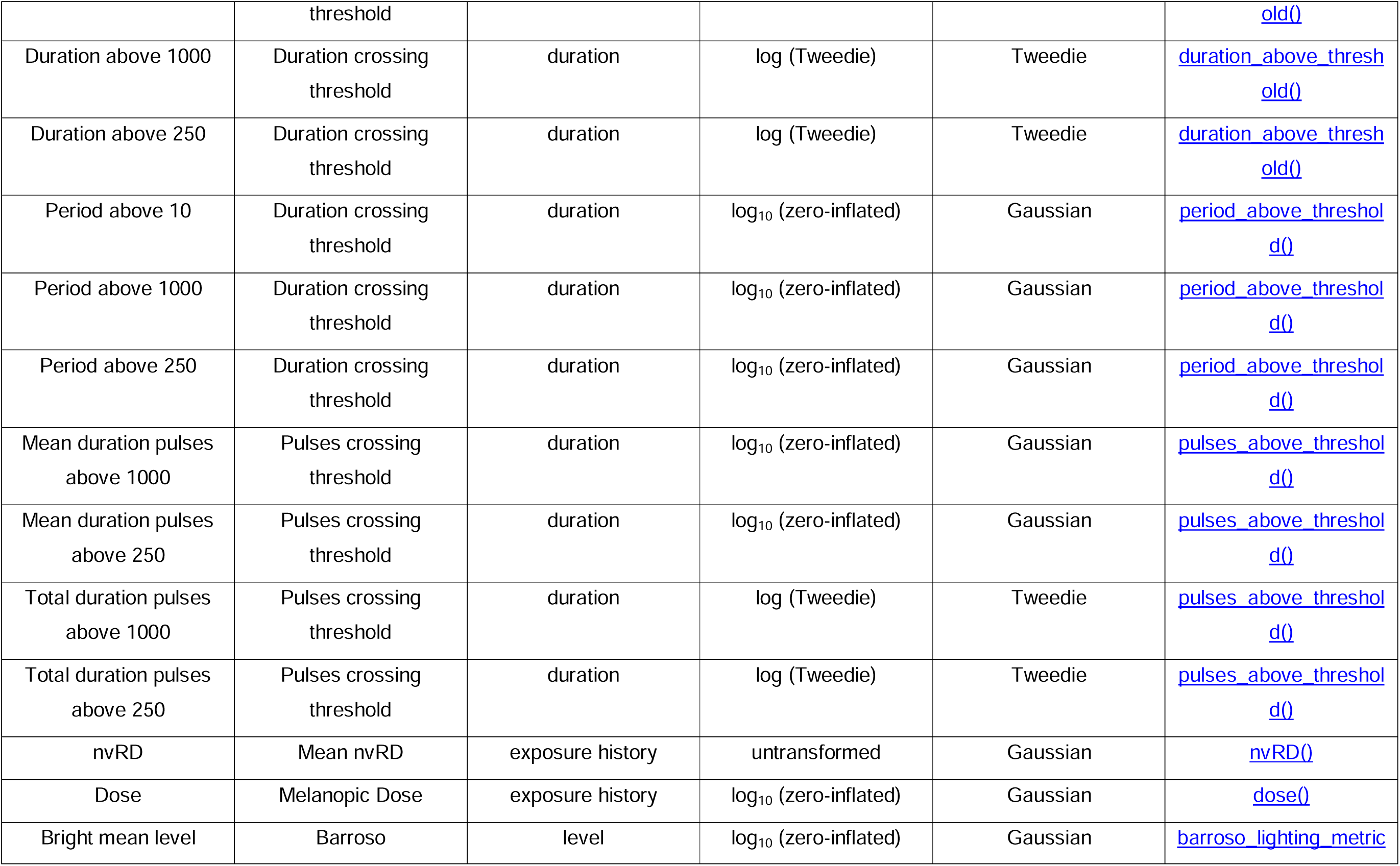
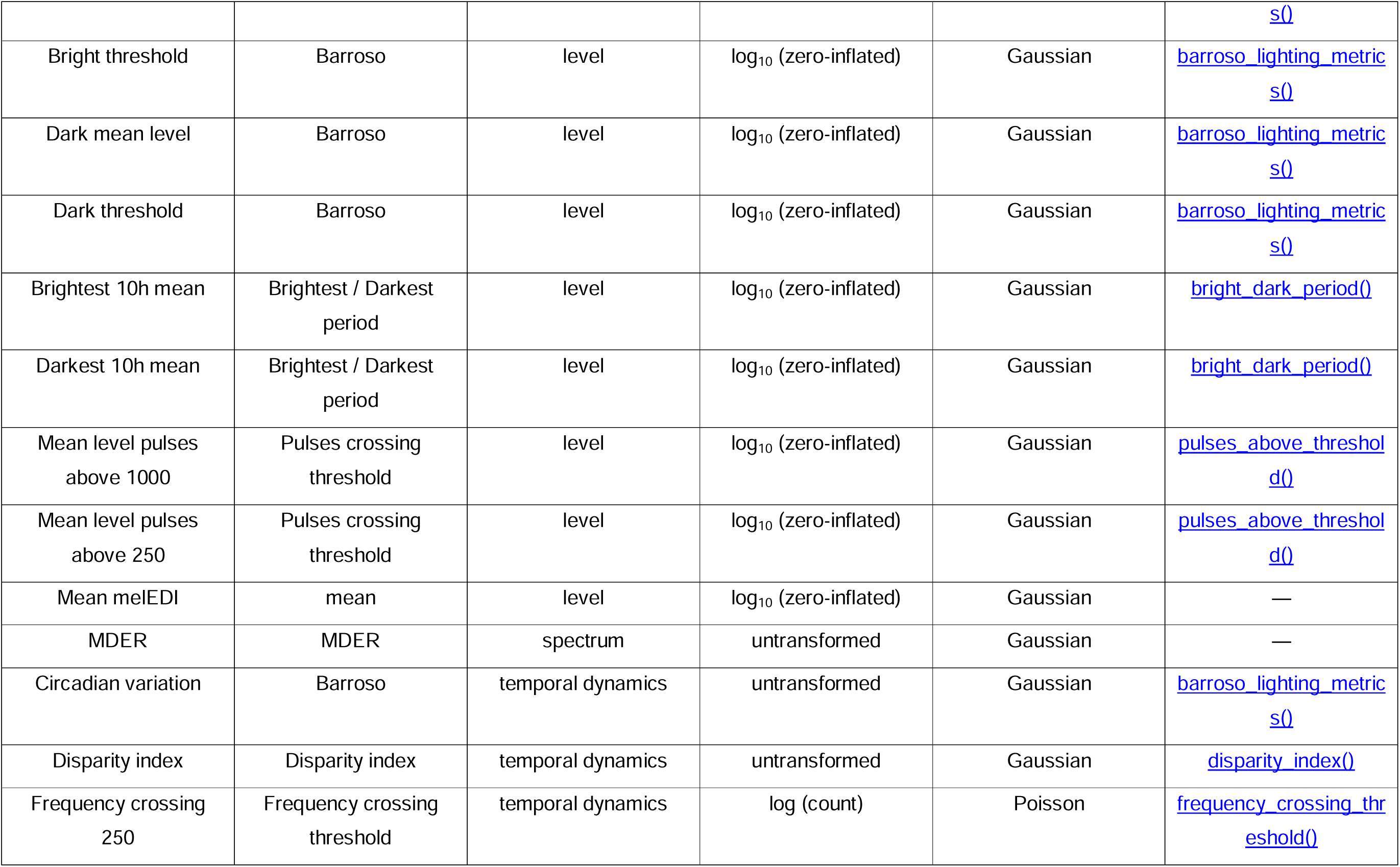
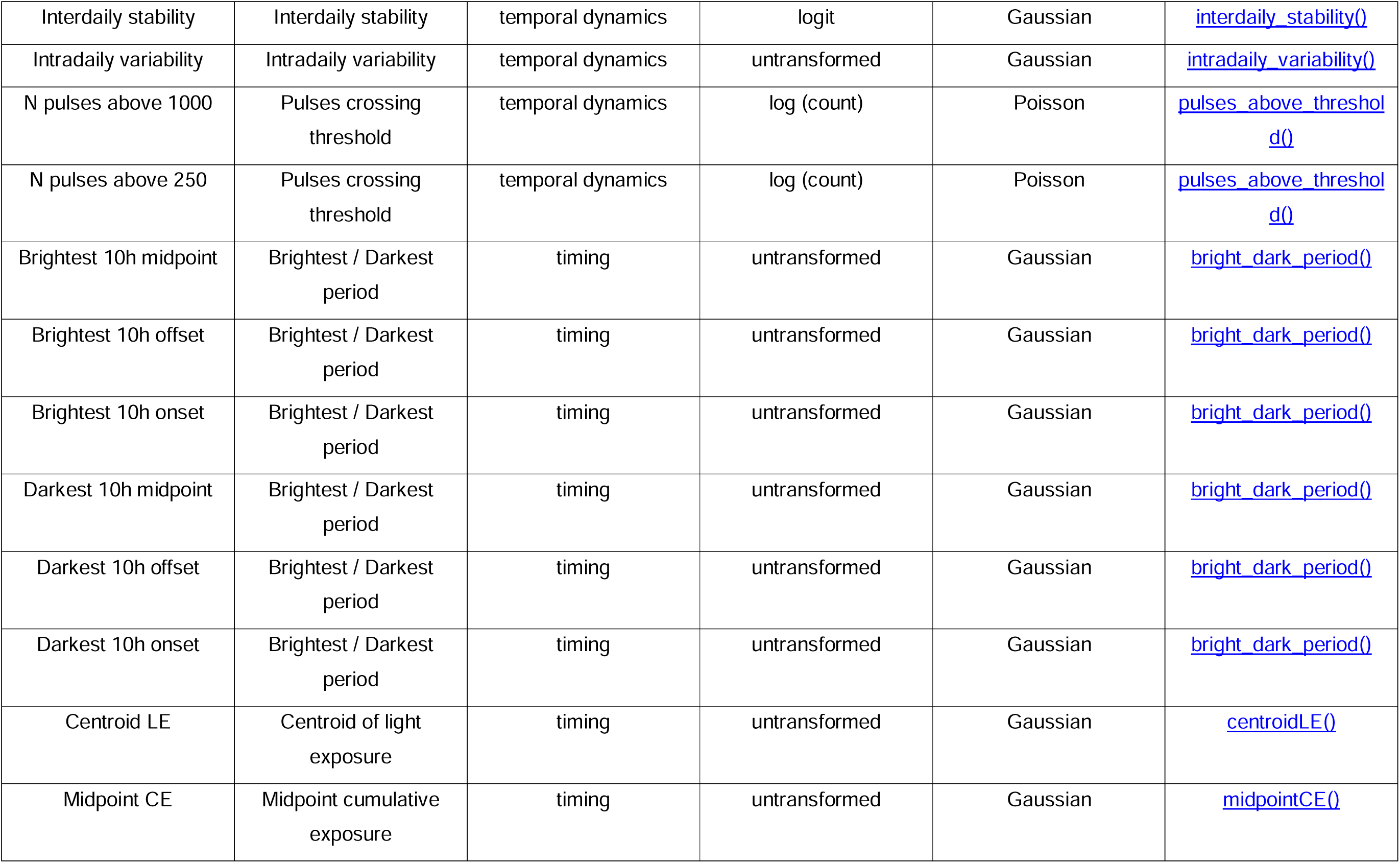
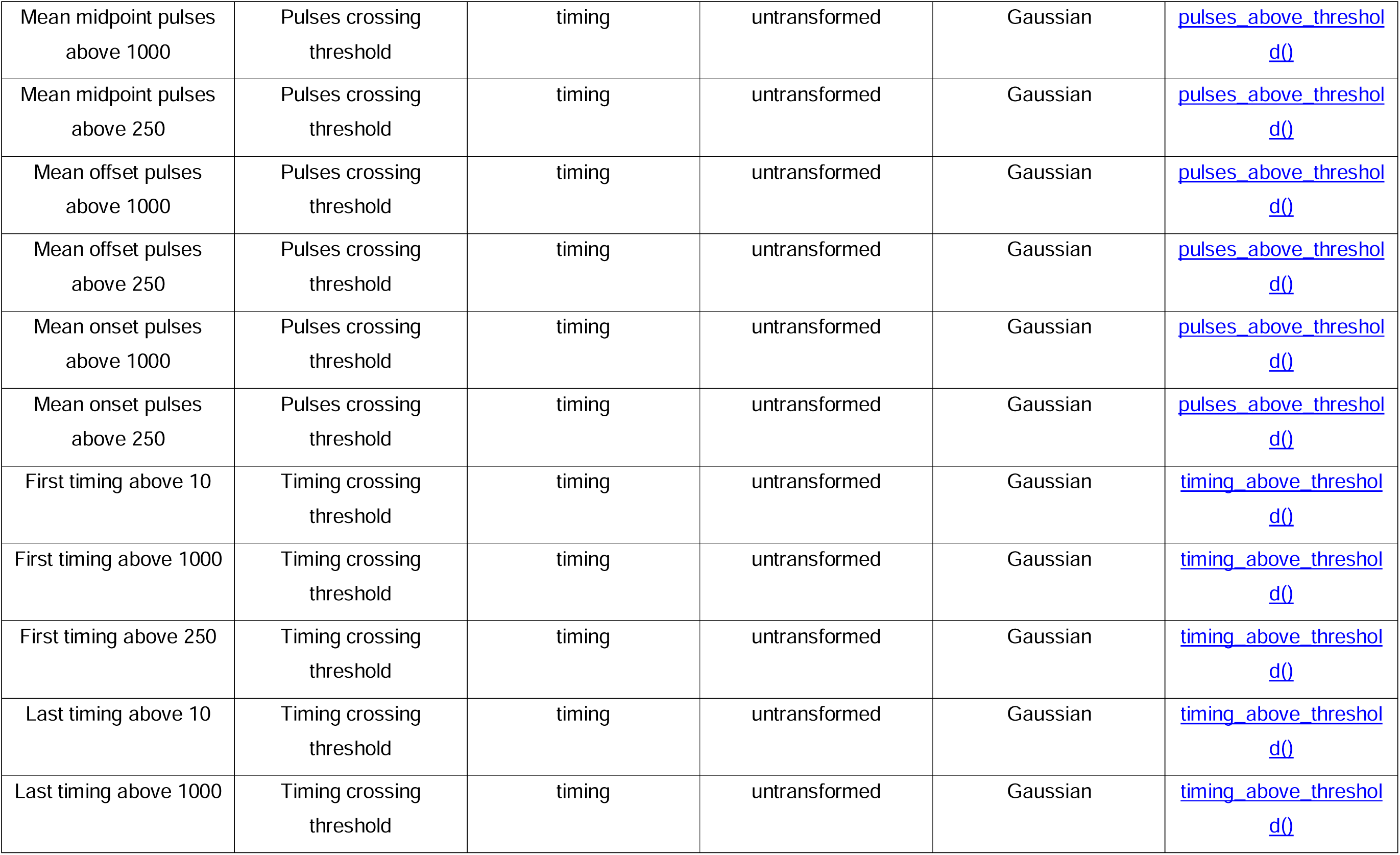
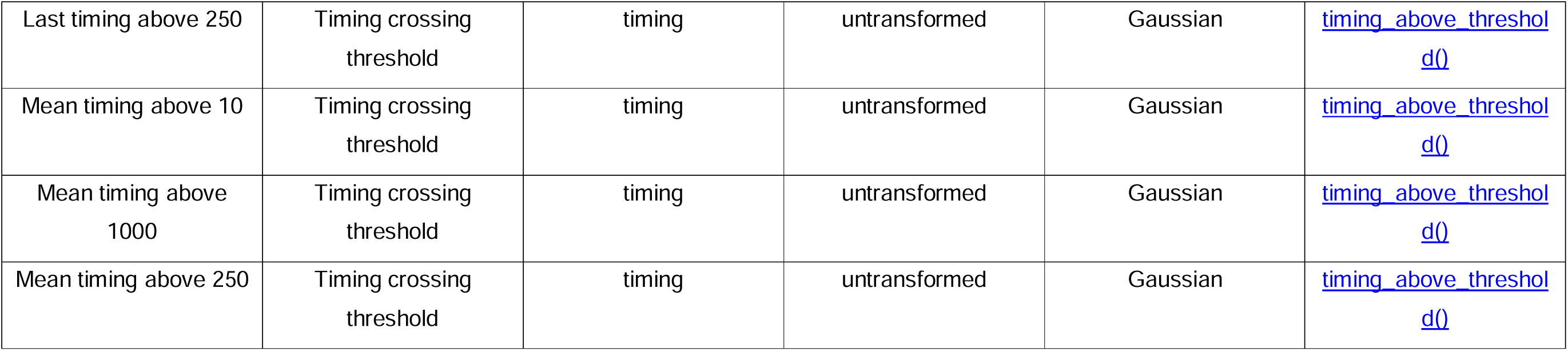
Modelling specification for all 54 daily light-exposure metrics. Scale indicates the transformation applied to the response variable before or during modelling. Distribution denotes the distributional family used in the mixed-effects model. All models used the same fixed-effects structure (site × position); see Equation 3 for the full formula. The LightLogR function column links to the LightLogR R package^36^ function used to compute each metric; — indicates metrics not computed by a dedicated package function.

**Supplementary Table S3:** Four sets of contextual classifications used to group placement errors by measurement context. Each set was defined as a combination of data from the contextual reports, photoperiod, and/or local time of day. The sets represent overlapping classifications of the same measurements rather than mutually exclusive partitions. ^1^Time of day was modelled as a cyclic smooth; the 15-minute discretisation was used only to extract estimates, not in model fitting.

| Set of contextual classifications | Categories | Number of unique classifications |
| --- | --- | --- |
| Environment & photoperiod | (indoors or outdoors) × (day or night) | 4 |
| Environment & time of day | (indoors or outdoors) × (15-minute interval<sup>1</sup>) | 192 |
| Activity & photoperiod | (awake at home or working in the office/home or in a vehicle or time outdoors) × (day or night) | 8 |
| Indoor work lighting | (electric or daylight or display) | 3 |

**Supplementary Table S4:**
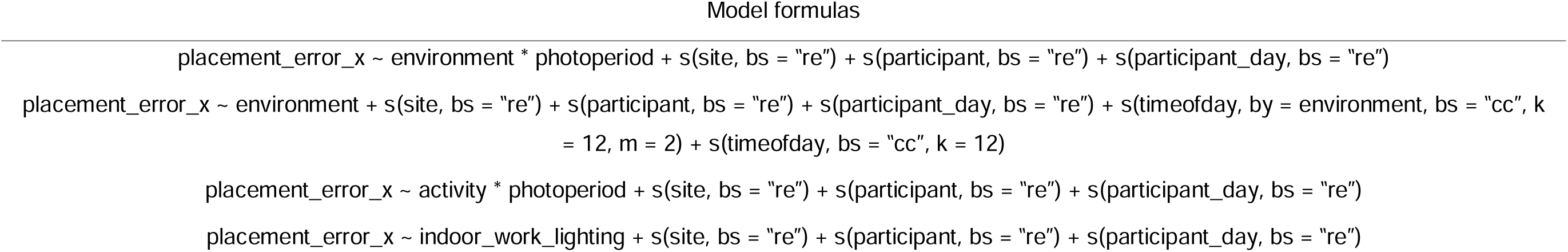
Model formulas for the eight fitted generalised additive mixed models (GAMMs). The subscript x denotes the placement (chest or wrist), with each formula fitted separately for both placements. Participants were implicitly nested within sites since each participant belonged to only one site and had a unique identifier. Because measurement days were not unique across participants, a constructed grouping factor (participant_day) was used to nest days within participants.

**Supplementary Table S5.**
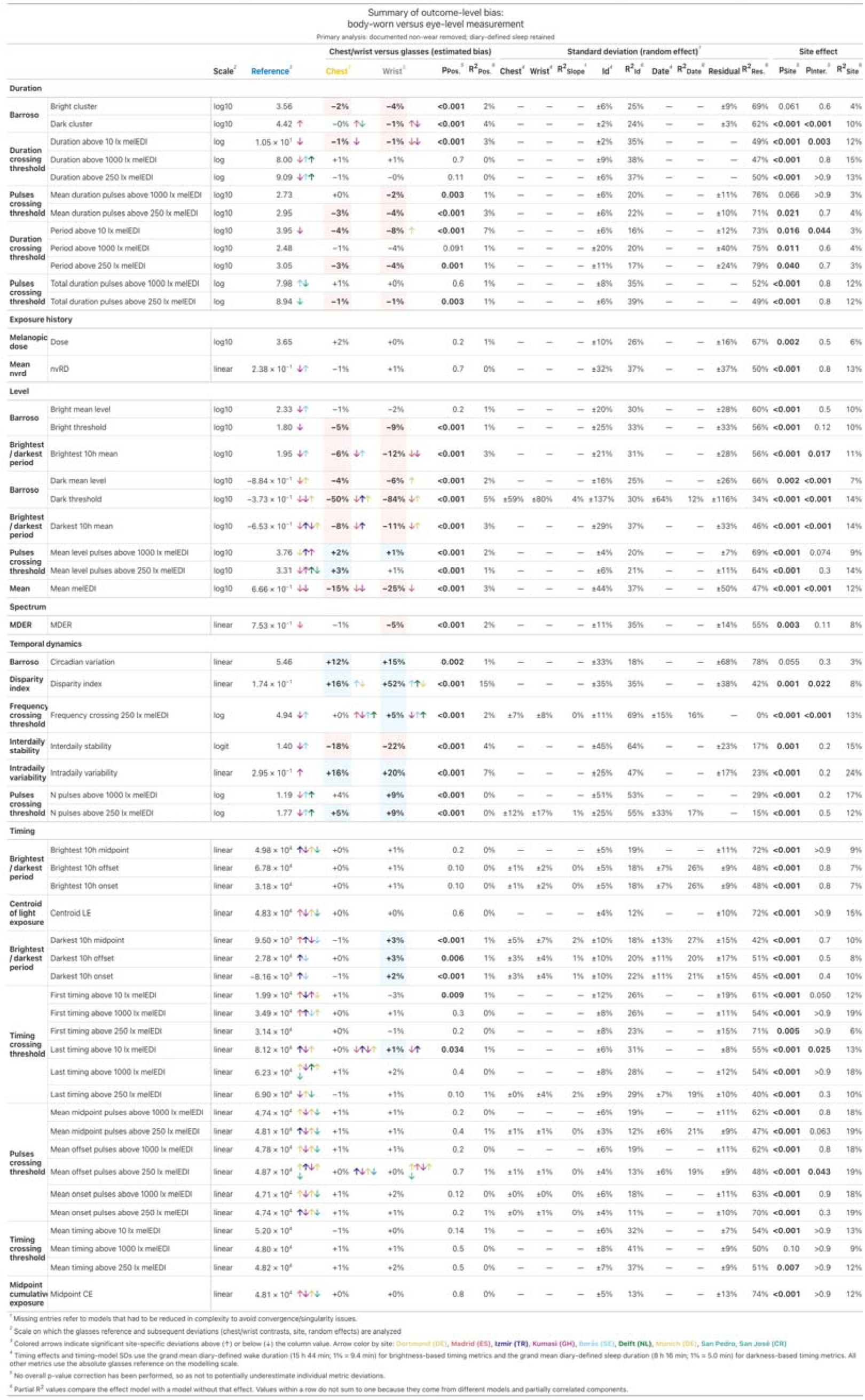

**Supplementary Table S6:** Sensitivity reproduction of the main outcome-level metric- summary table using the adjusted retained-non-wear dataset, in which documented non-wear and diary-defined sleep were both retained.

| Summary overview of outcome-level bias<br>for body-worn versus eye-level measurement |  |  |  |
| --- | --- | --- | --- |
| Sensitivity analysis: documented non-wear retained; diary-defined sleep retained<br>Timing metrics use mean wake/sleep durations;<br>other metrics to the absolute glasses reference. |  |  |  |
|  | Chest<sup>1</sup> | Wrist<sup>1</sup> | General<sup>1,2</sup> |
| Duration | bias: 1% (0% - 4%)<br>signif: 4/12, direction: +4/-8 | bias: 1% (0% - 8%)<br>signif: 6/12, direction: +3/-9 | R<sup>2</sup><sub>Pos.</sub>: 2% (1%-7%)<br>R<sup>2</sup><sub>Id</sub>: 25% (15%-37%)<br>SD<sub>Id</sub>: ±6% (2-20%) |
| Exposure history | bias: 1% (0% - 2%)<br>signif: 0/2, direction: +2/-0 | bias: 2% (1% - 3%)<br>signif: 0/2, direction: +2/-0 | R<sup>2</sup><sub>Pos.</sub>: ---<br>R<sup>2</sup><sub>Id</sub>: 30% (25%-36%)<br>SD<sub>Id</sub>: ±21% (10-31%) |
| Level | bias: 3% (0% - 19%)<br>signif: 5/9, direction: +3/-6 | bias: 7% (1% - 33%)<br>signif: 7/9, direction: +2/-7 | R<sup>2</sup><sub>Pos.</sub>: 2% (1%-4%)<br>R<sup>2</sup><sub>Id</sub>: 30% (20%-36%)<br>SD<sub>Id</sub>: ±22% (4-71%) |
| Spectrum | bias: 1% (1% - 1%)<br>signif: 0/1, direction: +0/-1 | bias: 5% (5% - 5%)<br>signif: 1/1, direction: +0/-1 | R<sup>2</sup><sub>Pos.</sub>: 2% (2%-2%)<br>R<sup>2</sup><sub>Id</sub>: 36% (36%-36%)<br>SD<sub>Id</sub>: ±11% (11-11%) |
| Temporal dynamics | bias: 6% (1% - 18%)<br>signif: 4/7, direction: +6/-1 | bias: 11% (6% - 57%)<br>signif: 7/7, direction: +6/-1 | R<sup>2</sup><sub>Pos.</sub>: 2% (1%-16%)<br>R<sup>2</sup><sub>Id</sub>: 53% (19%-68%)<br>SD<sub>Id</sub>: ±32% (11-53%) |
| Timing | bias: 0% (0% - 2%)<br>signif: 1/23, direction: +20/-3 | bias: 1% (0% - 4%)<br>signif: 7/23, direction: +21/-2 | R<sup>2</sup><sub>Pos.</sub>: 1% (0%-1%)<br>R<sup>2</sup><sub>Id</sub>: 19% (12%-28%)<br>SD<sub>Id</sub>: ±6% (4-12%) |
| Overall | bias: 1% (0% - 19%)<br>signif: 14/54, direction: +35/-19 | bias: 2% (0% - 57%)<br>signif: 28/54, direction: +34/-20 | R<sup>2</sup><sub>Pos.</sub>: 1% (0%-16%)<br>R<sup>2</sup><sub>Id</sub>: 23% (12%-68%)<br>SD<sub>Id</sub>: ±8% (2-71%) |
| Overall (significant) | bias: 5% (0% - 19%)<br>direction: +7/-7 | bias: 4% (1% - 57%)<br>direction: +13/-15 | R<sup>2</sup><sub>Pos.</sub>: 1% (0%-16%)<br>R<sup>2</sup><sub>Id</sub>: 23% (12%-68%)<br>SD<sub>Id</sub>: ±8% (2-71%) |

median bias (min - max), significant bias / all metrics, positive / negative deviations
<sup>1</sup> Timing effects and timing-model SDs use the grand mean diary-defined wake duration (15 h 45 min; 1% = 9.4 min) for brightness-based timing metrics and the grand mean diary-defined sleep duration (8 h 15 min; 1% = 5.0 min) for darkness-based timing metrics. All other metrics use the absolute glasses reference on the modelling scale. Overall rows combine these normalization bases.
<sup>2</sup> R<sup>2</sup><sub>Pos.</sub>: partial R<sup>2</sup> for the specific position effect (glasses, chest, or wrist; median min-max), significant metrics only. R<sup>2</sup><sub>Id</sub> and SD<sub>Id</sub>: individual ID random effect, all metrics.

**Supplementary Table S7:** Sensitivity reproduction of the main outcome-level metric-summary table using the adjusted wake-only dataset, in which documented non-wear and diary-defined sleep were removed before metric calculation. The table contains the 42 metrics available under this state definition.

| Summary overview of outcome-level bias<br>for body-worn versus eye-level measurement |  |  |  |
| --- | --- | --- | --- |
| Sensitivity analysis: documented non-wear and diary-defined sleep removed<br>Timing metrics use mean wake/sleep durations;<br>other metrics to the absolute glasses reference. |  |  |  |
|  | Chest<sup>1</sup> | Wrist<sup>1</sup> | General<sup>1,2</sup> |
| Duration | bias: 1% (0% - 4%)<br>signif: 4/11, direction: +3/-8 | bias: 2% (0% - 8%)<br>signif: 6/11, direction: +2/-9 | R<sup>2</sup><sub>Pos.</sub>: 3% (1%-6%)<br>R<sup>2</sup><sub>Id</sub>: 23% (16%-38%)<br>SD<sub>Id</sub>: ±6% (2-20%) |
| Exposure history | bias: 2% (2% - 2%)<br>signif: 0/1, direction: +1/-0 | bias: 0% (0% - 0%)<br>signif: 0/1, direction: +1/-0 | R<sup>2</sup><sub>Pos.</sub>: ---<br>R<sup>2</sup><sub>Id</sub>: 26% (26%-26%)<br>SD<sub>Id</sub>: ±10% (10-10%) |
| Level | bias: 4% (1% - 10%)<br>signif: 5/6, direction: +2/-4 | bias: 6% (1% - 16%)<br>signif: 4/6, direction: +2/-4 | R<sup>2</sup><sub>Pos.</sub>: 2% (1%-3%)<br>R<sup>2</sup><sub>Id</sub>: 29% (20%-33%)<br>SD<sub>Id</sub>: ±18% (4-27%) |
| Spectrum | bias: 1% (1% - 1%)<br>signif: 0/1, direction: +0/-1 | bias: 4% (4% - 4%)<br>signif: 1/1, direction: +0/-1 | R<sup>2</sup><sub>Pos.</sub>: 2% (2%-2%)<br>R<sup>2</sup><sub>Id</sub>: 34% (34%-34%)<br>SD<sub>Id</sub>: ±11% (11-11%) |
| Temporal dynamics | bias: 4% (1% - 5%)<br>signif: 1/3, direction: +3/-0 | bias: 9% (5% - 9%)<br>signif: 3/3, direction: +3/-0 | R<sup>2</sup><sub>Pos.</sub>: 0% (0%-2%)<br>R<sup>2</sup><sub>Id</sub>: 55% (53%-68%)<br>SD<sub>Id</sub>: ±26% (12-51%) |
| Timing | bias: 1% (0% - 2%)<br>signif: 1/20, direction: +16/-4 | bias: 1% (0% - 2%)<br>signif: 1/20, direction: +13/-7 | R<sup>2</sup><sub>Pos.</sub>: 1% (0%-1%)<br>R<sup>2</sup><sub>Id</sub>: 19% (11%-28%)<br>SD<sub>Id</sub>: ±6% (4-12%) |
| Overall | bias: 1% (0% - 10%)<br>signif: 11/42, direction: +25/-17 | bias: 1% (0% - 16%)<br>signif: 15/42, direction: +21/-21 | R<sup>2</sup><sub>Pos.</sub>: 2% (0%-6%)<br>R<sup>2</sup><sub>Id</sub>: 22% (11%-68%)<br>SD<sub>Id</sub>: ±7% (2-51%) |
| Overall (significant) | bias: 3% (1% - 10%)<br>direction: +4/-7 | bias: 4% (1% - 16%)<br>direction: +5/-10 | R<sup>2</sup><sub>Pos.</sub>: 2% (0%-6%)<br>R<sup>2</sup><sub>Id</sub>: 22% (11%-68%)<br>SD<sub>Id</sub>: ±7% (2-51%) |

median bias (min - max), significant bias / all metrics, positive / negative deviations
<sup>1</sup> Timing effects and timing-model SDs use the grand mean diary-defined wake duration (15 h 44 min; 1% = 9.4 min) for brightness-based timing metrics and the grand mean diary-defined sleep duration (8 h 16 min; 1% = 5.0 min) for darkness-based timing metrics. All other metrics use the absolute glasses reference on the modelling scale. Overall rows combine these normalization bases.
<sup>2</sup> R<sup>2</sup><sub>Pos.</sub>: partial R<sup>2</sup> for the specific position effect (glasses, chest, or wrist; median min-max), significant metrics only. R<sup>2</sup><sub>Id</sub> and SD<sub>Id</sub>: individual ID random effect, all metrics.

**Supplementary Table S8:** First sample size at which the pointwise metric-class median bias SD reached 5%, evaluated for 1–200 participants with three or seven days each and 1–7 days per participant with 25 or 50 participants, using 10,000 bootstrap replicates per design. The analysis included all 52 metrics with participant-day estimates; IS and IV were excluded because daily estimates were unavailable.

| Sample size for class-median bias $SD \leq 5\%$ | | | | | | | | | |
| --- | --- | --- | --- | --- | --- | --- | --- | --- | --- |
| 52 daily metrics; pointwise median within class; B = 10,000 bootstrap draws |  |  |  |  |  |  |  |  |  |
| Metric class | Metrics | N participants |  |  |  | Days per participant |  |  |  |
|  |  | 3 days each |  | 7 days each |  | 25 participants |  | 50 participants |  |
|  |  | Chest | Wrist | Chest | Wrist | Chest | Wrist | Chest | Wrist |
| Duration | 12 | 3 | 4 | 3 | 3 | 1 | 1 | 1 | 1 |
| Exposure history | 2 | 7 | 10 | 5 | 8 | 1 | 1 | 1 | 1 |
| Level | 9 | 10 | 21 | 7 | 16 | 1 | 2 | 1 | 1 |
| Spectrum | 1 | 3 | 5 | 2 | 4 | 1 | 1 | 1 | 1 |
| Temporal dynamics | 5 | 45 | 76 | 33 | 59 | — | — | 3 | — |
| Timing | 23 | 3 | 4 | 3 | 3 | 1 | 1 | 1 | 1 |

**Note.** Cells give the first N participants (left) or days per participant (right) meeting the tolerance. — indicates no crossing within the evaluated range. A class-median crossing does not imply that every metric in the class meets the tolerance.

